# *Drosophila* germ band extension: a two-state reshaping mechanism

**DOI:** 10.64898/2026.03.09.710653

**Authors:** Tianyi Zhu, Hongkang Zhu, Ben O’Shaughnessy

## Abstract

During embryogenesis tissues are reshaped by diverse forces that elicit fluid-like or solid-like response, but the underlying principles that determine tissue state are unclear. Here we study *Drosophila* germ-band extension (GBE), a classic example of convergent-extension that extends the ventral germ band (GB) to the dorsal embryo surface. Using experimentally constrained biophysical modeling we find fluctuating internal cell-cell junctional myosin stresses excite cell intercalations that weakly perturb the anterior GB tissue into a fluidized state. Planar-polarized intercalation bias drives weak convergent-extension of the fluid. In the posterior GB, by contrast, external stress from adjacent tissue induces crystal-like ordering of the cells. External stress-induced cell intercalations mediate crystal defect annealing and plastic flow of the solid that wraps the GB around the narrow embryo posterior. Thus, the inhomogeneous reshaping challenge of GBE is dealt with by a two-state remodeling strategy in which a slowly remodeled anterior fluid coexists with a posterior solid undergoing fast plastic flow. Similar strategies find use in other contexts such as wound healing, where actomyosin boundary stresses drive crystal-like order that imposes shape regularity on the inner wound boundary for successful wound closure.

## Introduction

Embryogenesis transforms a single-celled zygote into a fully developed embryo with germ layers, an established body plan, and rudimentary organs. In accordance with this program, tissues undergo specific reshaping episodes driven by many forces including actomyosin contractility, cell-cell adhesion, proliferative forces, and viscous and elastic forces (1-4). These forces drive tissue shape changes that depend on the mechanical response characteristics of the tissue, fluid-like or solid-like, with interesting parallels to inanimate materials (5-7). Both fluid-like and solid-like tissue responses are seen in basic morphogenetic processes such as body axis elongation (8, 9), epiboly (10, 11) and branching morphogenesis (12).

A widely used mode of tissue reshaping is convergent-extension, a conserved mechanism that extends tissue in one direction while contracting it in the perpendicular direction (13, 14). In vertebrates, convergent-extension is associated with mediolateral cell intercalation regulated by the planar cell polarity (PCP) signaling pathway (15). Vertebrates use convergent-extension during neurulation to elongate and narrow the neural plate in the anterior-posterior (AP) and mediolateral directions (16), respectively, while other extensively studied examples include notochord extension in amphibians (17) and primitive streak extension in chick (18).

The classic example of convergent-extension in invertebrates is body axis elongation in the *Drosophila* embryo, driven by germ band extension (GBE). GBE establishes the segmental embryo organization and coordinates with posterior midgut (PMG) invagination to incorporate the primordial germ cells known as pole cells (19, 20). The germ band (GB) is an epithelial sheet on the ventral and lateral surface of the embryo trunk which GBE extends along the AP axis and narrows in the ventral-lateral (VL) direction. The tissue reshaping is facilitated by mediolateral cell intercalations associated with myosin planar polarity regulated by pair-rule genes and downstream Toll family receptors (21, 22), with enriched myosin II at VL-oriented cell-cell junctions extending across the GB in cable-like formations (21, 23). Additionally, Bazooka/Par-3 (24, 25) regulates enrichment of adherens junctions at AP-oriented cell-cell contacts (26, 27).

GBE is thought driven by the contractile tensions at these myosin-enriched cell-cell junctions. Large temporal fluctuations in myosin levels are measured (28), suggesting the tensions have similarly large fluctuations. Tensions are also highly spatially inhomogeneous: the anterior GB contains myosin cables (21), but junctional myosin is virtually absent from the posterior GB (29). While myosin stresses are presumably lower in the posterior, boundary stresses act on the posterior end of the GB due to concurrent invagination of the PMG primordium in the neighboring tissue (30-32). These boundary stresses appear to contribute to elongation of cells and the GB as a whole since in embryos from *tor*^-/-^ mothers where PMG invagination is abolished the GB is shorter, cells are less elongated and aligned, and new junctions generated by intercalation are less aligned parallel to the AP axis (30).

Does the GB respond to these mechanical stresses as a fluid or a solid? During the initial fast phase of GBE cell intercalation rates are high (33-36), suggesting the response is fluid-like. Specifically, the ∼ 1/20 min^−1^ intercalation rates per cell in the anterior GB (33, 36) exceed the ∼ 1/30 min^−1^strain rates (30), suggesting intercalative cell rearrangements are frequent enough to sustain fluid-like convergent-extension without cell deformation. Indeed, cell deformations are small in the anterior GB (30-32, 36), consistent with a fluid. However, despite high overall rates, intercalation rates are much lower in the posterior GB (34) and posterior cells undergo much larger AP elongations (30-32), features that suggest solid-like response.

Thus, the driving stresses and the tissue states are inhomogeneous throughout the GB. It is therefore unsurprising that the tissue deformation rates are also inhomogeneous. While *Drosophila* GBE is a classic example of convergent-extension, the reshaping is more complex than simple rectangle-to-rectangle elongation-contraction. During the rapid phase GBE wraps the GB around the posterior embryo pole and extends it onto the dorsal surface, an essential step to ensure morphological segments later produce distinct tissues according to the segmented body plan (37, 38). Thus, a geometric necessity is that the posterior GB be significantly narrowed to accommodate the high curvature of the posterior pole. To achieve this, convergence and extension are highly inhomogeneous: the posterior GB is severely narrowed and elongated, with only moderate deformation of the anterior portion of the GB.

Vertex models have been widely used to computationally simulate confluent tissue (30, 33, 39-46). Invoking areal elasticity and a perimeter elasticity that represents cortical tension and adhesion effects, these models predict fluid-like and solid-like tissue states, with a solid-to-fluid transition controlled by a shape index parameter, the ratio of the energetically preferred cell perimeter to the square root of preferred cell area (39, 43). The framework is suited to modeling GBE, but a realistic description must account for the fluctuating, planar-polarized and spatially inhomogeneous junctional tensions. A vertex model predicted that anisotropic tensions promote tissue solidification by aligning cells (33). In another vertex model study, temporal fluctuations in junctional tensions promoted tissue fluidization, but solid-like behavior was recovered when the fluctuation persistence times were sufficiently long (47). Mechanosensitive effects were invoked in another study, leading to intercalations via a positive feedback mechanism in which high tension junctions further recruited myosin at the expense of neighboring junctions (48, 49). These works revealed fundamental features of tissue mechanics. However, to our knowledge the combination of conditions during GBE has not been addressed, where planar polarized junctional tensions have cable patterning and strong temporal fluctuations, and conditions in the anterior and posterior GB are very different.

Here we studied GBE in a vertex model framework, with experimentally set junctional myosin distributions and posterior boundary forces from PMG invagination. We find the reshaping task required of GBE is accomplished by activating distinct tissue states in the anterior and posterior GB. The anterior tissue is internally fluidized by junctional myosin fluctuations, activating a weakly perturbed fluid state where anisotropic cell intercalations mediate continuous convergent-extension deformation without structural change at the cellular level. By contrast, the posterior GB is solidified by external stresses that severely stretch cells and catalyze their organization into crystal-like tissue domains. Stress-induced intercalations promote plastic flow, and mediate crystal defect annealing and remodeling of the solid-like posterior tissue, facilitating its dramatic elongation and narrowing to wrap around the embryo posterior. Thus, tissue reshaping during GBE is accomplished by co-activation of fluid and solid tissue states across the GB.

## Results

### Model

The GB is a sector of epithelial cells on the ventral and lateral surface of the early *Drosophila* embryo trunk that later contributes to the ectoderm, one of the three germ layers (24, 50-52). GBE is a major morphogenetic event (53, 54), a classic example of convergent-extension deformation that reshapes the GB to almost twice its original length, extending the posterior GB around the embryo posterior onto the dorsal surface (55) (Fig. 1A). Extension to the dorsal surface is required for the process of segmentation (37, 56, 57) which creates the body plan. Lasting ∼ 2 hrs, GBE commences ∼ 3 hr 15 min after fertilization when the *Drosophila* embryo consists of ∼ 6000 cells. Our model addresses the initial ∼ 30 min rapid phase when most morphological changes occur (55). The GB measures ∼ 50 cells in the AP direction and ∼ 40 cells in the VL direction, its lateral edges connecting to the extraembryonic amnioserosa epithelial tissue on the dorsal surface (55, 57, 58) so that GBE stretches the amnioserosa cells in the VL and AP directions (Figs. 1A, B). The anterior GB connects to the cephalic furrow (55, 59) whose development by tissue invagination continues through GBE, while the posterior GB edge connects to the invaginating PMG primordium.

**Figure 1.**
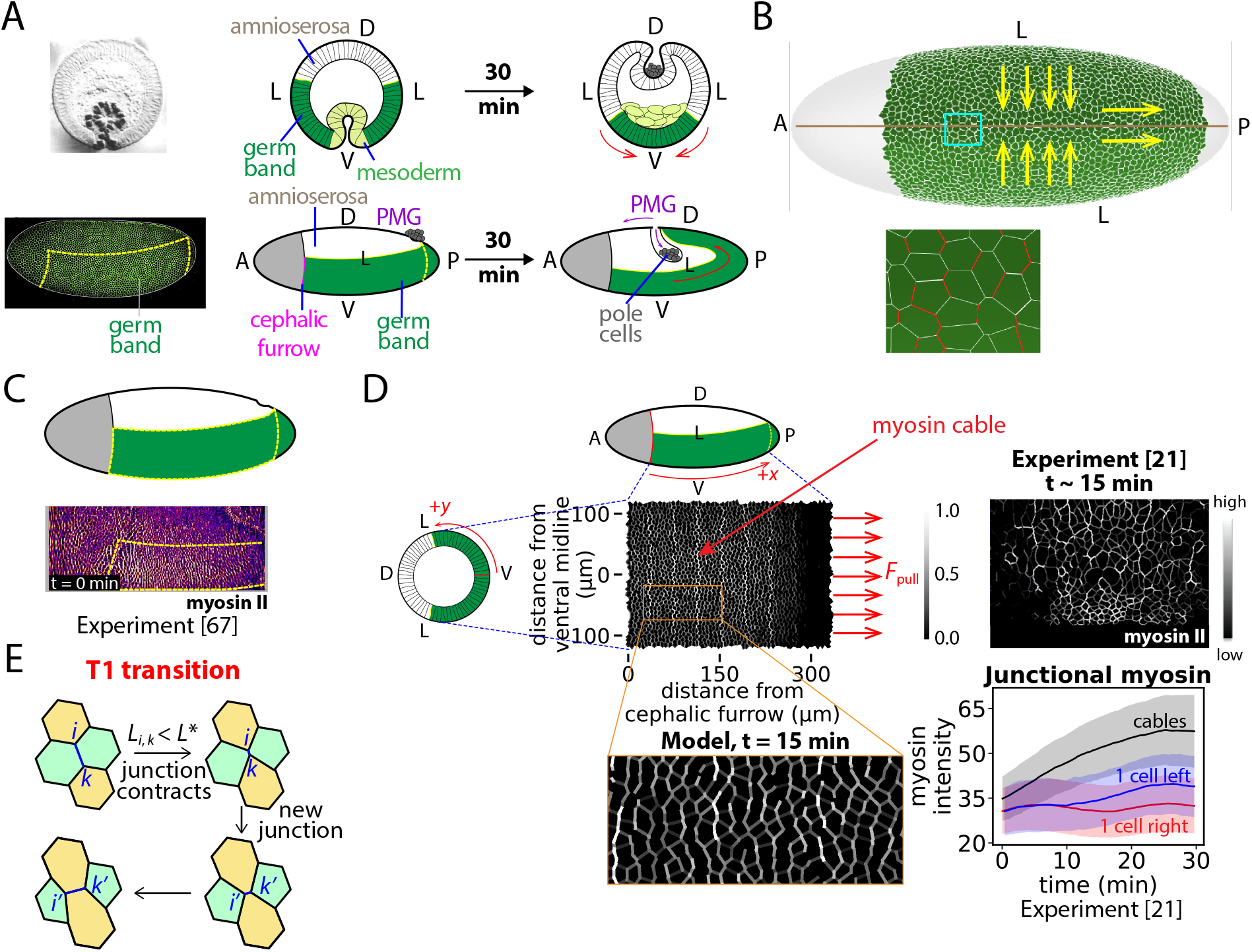
Model of *Drosophila* germ-band extension (GBE). (A) At GBE onset the germ band (GB) extends from the cephalic furrow to the embryo posterior on the ventral (V) surface (dashed lines). Over 30 min (GBE fast phase) it narrows and elongates in the ventral-lateral (VL) and anterior-posterior (AP) directions, respectively, extending to the dorsal surface (D). The GB connects to the posterior midgut (PMG) primordium whose invagination pulls its rear boundary, assisting GBE. Top row: cross-sections. Bottom row: lateral views. Images at GBE onset, adapted from ref. (57) (top left) and ref. (23) (bottom left). (B) Schematic of epithelial cellular network in the GB (ventral view). AP-oriented cell-cell junctions are myosin-enriched (red). GBE is a convergent-extension reshaping (yellow arrows). (C) Myosin levels are high and low in the anterior and posterior GBs, respectively. Light sheet microscopy image adapted from ref. (67), world-map 2D projection. (D) Model setup. Initially the GB extends ∼ 50 cells in the AP direction from the cephalic furrow (*x* = 0) and ∼ 40 cells in the VL direction symmetric about the ventral midline (*y* = 0). GBE is driven by junctional myosin (white) enriched in VL-oriented cables in the anterior GB and posterior boundary forces from PMG invagination (arrows). For comparison, microscopy image cropped from ref. (21) shows myosin cables in a region from ventral midline to lateral embryo edge. Myosin levels for cable and neighboring junctions during GBE fast phase, replotted from ref. (21) (bottom right). Shaded regions, 95% confidence intervals. (E) A T1 transition facilitating cell intercalation is triggered when a junction becomes shorter than a threshold *L*^∗^.

Here we develop a biophysical model of the rapid phase of *Drosophila* GBE. To describe the epithelial GB cells on the ventral embryo surface we use a vertex model which is then mapped onto the 3D embryo surface approximated as the surface of an ellipsoid with principal axes of lengths 490 μm, 180 μm and 180 μm taken from experimentally reported embryo dimensions (49). The force balance on vertex *i* reads

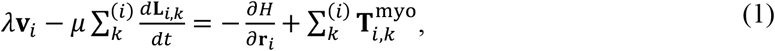

where the vector **L**_*i,k*_ represents the cell-cell junction from vertex *i* to *k*, **r**_*i*_ and **v**_*i*_ are the position and velocity of vertex *i, μ* is the junctional viscosity, *λ* the external drag coefficient and 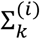 sums over all vertices *k* connected by a junction to vertex *i*. The level of excess actomyosin material at junction (*i, k*) determine the actomyosin tension 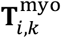, whose spatiotemporal distribution drives tissue remodeling (see below). The Hamiltonian *H* has the commonly used vertex model form (39, 44),

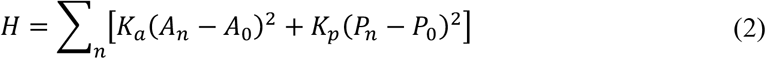

where *A*_*n*_ and *P*_*n*_ are the apical area and perimeter of cell *n, A*_0_ and *P*_0_ the preferred values, and the sum is over cells *n* in the GB. The areal elasticity constant *K*_*a*_ represents resistance to volume change (60), given cell heights remain approximately constant (61), while the perimeter elastic constant *K*_*p*_ quantifies the net effect of cell-cell adhesion and actomyosin junctional tension in the reference state (tending to increase and decrease perimeter length, respectively).

#### Cell intercalation

Cell intercalation by a T1 transition (30, 62) occurs spontaneously if a junction length drops below a threshold *L*^∗^ chosen to be 2% of the initial mean junction length. A new orthogonal junction of length 2*L*^∗^ is then born, Fig. 1E.

#### Experimental background: junctional myosin during GBE

GBE is driven in part by actomyosin tensions at apical cell-cell junctions (28, 62-64) (Fig. 1B). Experimentally, GB myosin ramps up during the last 5 min of ventral furrow formation that precedes GBE, the total amount thereafter remaining constant during the rapid GBE phase (21, 62). The activated myosin localizes primarily to cell-cell junctions (62, 65), organized in cables oriented in the VL direction and concentrated in the anterior GB. The posterior GB has much lower myosin content, Figs. 1C, D. More quantitatively: (1) In the VL direction junctional myosin decreases with distance from the ventral midline. In the AP direction, myosin is enriched and approximately constant in the anterior ∼ 60% of the GB, decaying to almost zero in the posterior GB, where the anterior/posterior boundary is fixed relative to the embryo and does not advect with the GB tissue (29, 66, 67). (2) Junctional myosin is planar polarized, with higher myosin at more VL-aligned junctions (21, 28, 62, 67). (3) Junctions with high activated myosin tend to form multicellular cables extending in the VL direction, about one cable every five VL cell rows (21), Fig. 1D. The cable forces during GBE are not established, but apical actomyosin networks exerted forces 1 − 1.5 *p*N during *Drosophila* dorsal closure (68). Non-cable myosin at other VL-oriented junctions has about half the myosin amount (21). (4) New junctions created by T1 transitions have much lower myosin (69), remaining low for at least 10 min (30).

#### Junctional actomyosin tensions in the model, 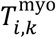

The model implements these experimental features (see Supplementary Information). One of every five (VL-oriented) cell rows carries a myosin cable with high junctional tensions, while non-cable myosin tensions are one half the cable levels: the mean tension at junction (*i, k*) is 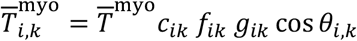, where 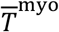 defines the characteristic junctional tension (Table 1), *c*_*ik*_ = 1 for cable junctions, and *c*_*ik*_ = 1/2 for non-cable junctions (Fig. 1D). Here *f*_*ik*_, *g*_*ik*_ are the envelope factors for the junction (*i, k*), related to the VL spatial envelope *f*(*y*) and the AP envelope *g*(*x*) where *x* and *y* are measured in the AP and VL directions, respectively. The VL envelope *f*(*y*) is bell-shaped (Fig. 1D and Supplementary Information), while the AP envelope *g*(*x*) equals unity in the anterior 60% of the GB, then decaying in a Gaussian manner in the posterior 40%. The anterior/posterior boundary is at a fixed location *x* = 190 μm, positioning it 190 μm from the cephalic furrow at *x* = 0 which defines the anterior end of the GB, Fig. 1D. The factor *f*_*ik*_ is the mean of the envelope *f*(*y*) evaluated at the centroids of the two cells sharing junction (*i, k*), and similarly for *g*_*ik*_. To account for junctional myosin planar polarity the tension depends on the junction orientational angle *θ*_*i,k*_ relative to the VL direction.

**Table 1.** Model parameter values.

| Parameter |  | Value |  |
| --- | --- | --- | --- |
| $\bar{T}^{\text{myo}}$ | Characteristic junctional myosin tension | 1.5 nN | (A) |
| $K_a$ | Areal elasticity | 12.2 pN $\mu\text{m}^{-3}$ | (B) |
| $K_p$ | Perimeter elasticity | 176 pN $\mu\text{m}^{-1}$ | (C) |
| $\lambda$ | External drag coefficient | 1 pN s $\text{nm}^{-1}$ | (D) |
| $\mu$ | Junctional viscosity | 36 pN s $\text{nm}^{-1}$ | Ref. (74) |
| $F_{\text{pull}}$ | Posterior pulling force | 1550 pN per cell | (E) |
| $A_0$ | Preferred cell area | 42 $\mu\text{m}^2$ | Ref. (30) |
| $P_0$ | Preferred cell perimeter | 24 $\mu\text{m}$ | |
| $p_0$ | Target cell shape index | 3.72 | (F) |
| $L^*$ | Threshold junction length to trigger T1 transition | 83 nm | |

(A) Forces 1 – 1.5 pN were measured during *Drosophila* dorsal closure (68).
(B) Best-fit of mean predicted cell area decrease after 10 min of GBE to experimental value from ref. (30) (Fig. S2A).
(C) Best-fit of predicted mean cell aspect ratio change after 30 min of GBE to experimental value from ref. (30) (Fig. S2B).
(D) Best fit to reproduce observed total number of T1 transitions in WT GB, ref. (34) (Fig. S3).
(E) Best fit to reproduce measured GB elongation rate of 8 $\mu\text{m min}^{-1}$ , ref. (30) (Fig. S1).
(F) With this value the initial GB before myosin ramp-up is solid-like, since $p_0 < p_0^*$ where $p_0^* = 3.81$ is the value at the rigidity transition (39, 71).

#### Myosin fluctuations

Relative myosin fluctuations of ∼ 30% with persistence time ∼ 200 sec were experimentally measured at individual junctions over time, and among junctions of the same class (21, 28). Accordingly, the model implements fluctuating time-dependent junctional tensions 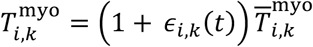 where 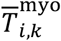 is the mean tension (see above) and *ϵ*_*i,k*_ (*t*) is a stochastic variable with zero mean, fluctuations of 0.3 and correlation time 200 sec. Temporal fluctuations are statistically independent across junctions. New junctions spawned by T1 intercalations are assumed tensionless for the remainder of the simulation, as experiments showed such junctions had virtually undetectable myosin (69) and reduced recoil velocity following laser ablation (30), and low myosin persisted for over 10 min (30).

#### Forces acting on GB boundaries

(1) To implement forces on the posterior boundary due to PMG invagination (30-32, 70) (Fig. 1A), the model applies forces that reproduce the boundary velocity of 8 μm min^−1^ measured in wild-type embryos (30, 53, 63). Following an initial ∼ 5 min transient, we find a constant required force of ∼ 1550 pN per cell (Fig. S1 and Table 1). (2) The lateral GB edges connect to the amnioserosa whose cells become highly elongated and irregularly shaped during GBE (34, 49). At the anterior GB edge lies the cephalic furrow, which bends during GBE as GB cells move in the anterior direction (32, 53). Since these observations suggest GB cells at the lateral and anterior edges can move relatively freely, the model assumes these boundaries are stress-free.

#### Initializing the GB

The GB is represented as a sector ∼ 40 cells wide (VL) and ∼ 50 cells long (AP). The Hamiltonian (Eq. (2)) is rewritten

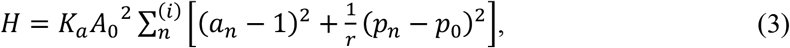

where 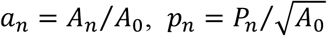 and *r* = (*K*_*a*_*A*_0_)*/K*_*p*_ is the inverse perimeter modulus. It was shown that a rigidity transition (39, 71) occurs at 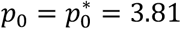, where 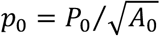 is the target cell shape index. Here, we choose *p*_0_= 3.72, so the GB lies in the solid-like regime before myosin activation, consistent with almost vanishing cell intercalation rates observed before and during the first 5 min of GBE (33).

The initial GB is generated by sequential random addition (SRA) of 1853 4μm radius disks to a rectangular domain followed by Voronoi tessellation. The disk centers will correspond to cell centroids. The lattice energy is then decreased following the dynamics of Eq. (1), but with zero actomyosin tensions 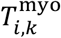 and zero junctional viscosity, together with T1 transitions. The procedure is halted when the energy reduction rate relative to the initial energy falls below 10^−9^ s^−1^.

#### Model parameters (Table 1)

The elasticities *K*_*a*_, *K*_9_, were chosen to reproduce the experimentally reported ∼ 10% mean cell area decrease and ∼ 20% cell aspect ratio decrease after 10 min of GBE (30) (Fig. S2). The external drag coefficient *λ* is likely dominated by friction between the translating GB cells and the vitelline membrane, the inner surface of the egg-shell (72). Its value was chosen to reproduce the experimental number of T1 transitions during GBE (34) (Fig. S3). The junctional viscosity *μ* describes dissipation due to anchoring of actomyosin structures to lateral cell membranes (73, 74) and other internal effects. We used the value estimated in ref. (74).

### Internal and external forces reshape the GB to wrap around the embryo posterior

*Drosophila* GBE is a classic example of convergent-extension (13, 75). However, the GB in fact undergoes a complex inhomogeneous reshaping, resulting in a relatively mildly deformed anterior GB but an extended, severely narrowed posterior GB that wraps around the high curvature embryo posterior and extends to the dorsal surface (49) (Fig. 1A). The inhomogeneity is presumably due to the distinct mechanical stress fields acting on different regions (see schematic of Fig. 2A): the anterior GB feels internal stresses from VL-oriented junctional myosin cables (21, 23), while the posterior GB experiences external force from adjacent tissue in the PMG primordium associated with PMG invagination (30-32). These features are incorporated into our model.

**Figure 2.**
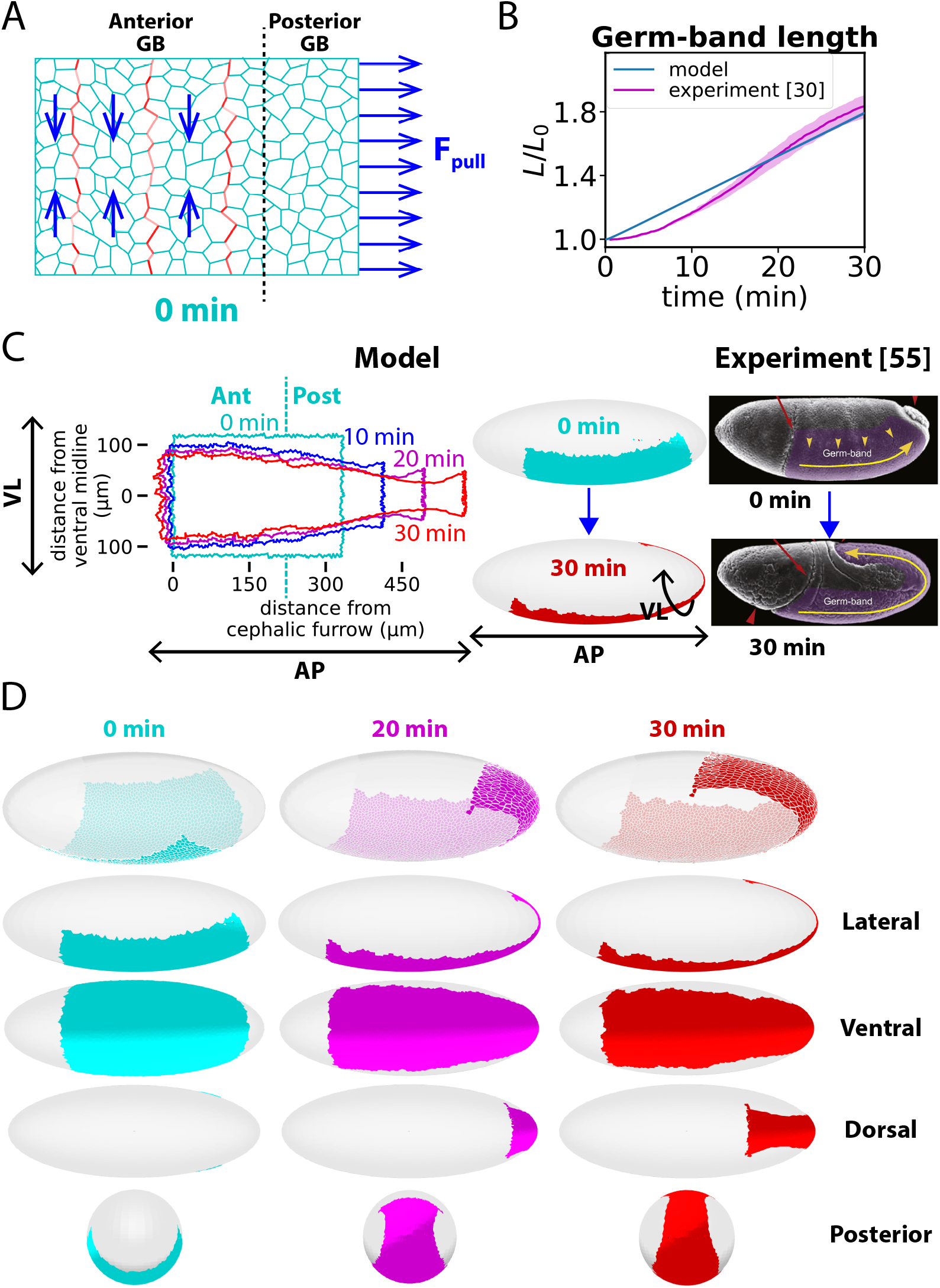
The model reproduces the experimental GB shape evolution. (A) Schematic illustrating inhomogeneous stresses driving GBE. Junctional actomyosin tensions act on the anterior GB from planar polarized myosin concentrated in cables (red). PMG invagination exerts pulling forces on the posterior GB boundary. (B) Model predicted relative GB length versus time during GBE (*n* = 10 embryos). Experimental data (*n* = 4 embryos) replotted from ref. (30). Shaded regions: SEM. (C) Predicted evolution of GB shape during GBE. VL, AP denote ventral-lateral, anterior-posterior directions, respectively. The simulated GB is mapped onto an ellipsoid closely approximating the *Drosophila* embryo surface (see model section). Right: Electron micrographs, adapted from ref. (55) (GB highlighted purple). (D) Predicted evolution of GB during GBE.

To study how this combination of stresses achieves the desired shape change, we simulated the 30-min rapid phase of GBE. Following initialization of the confluent GB cells, the junctional myosin is abruptly switched on at time zero to reflect the rapid ramp-up (21, 62) during the last 5 min of ventral furrow formation that precedes GBE. The model evolved the GB shape in quantitative agreement with experiment.

Over 30 min the GB elongated ∼ 1.8-fold (Fig. 2B), extending around the posterior end and onto the dorsal surface so the posterior GB boundary reached the mid-dorsal region (Figs. 2C, D). To wrap around the embryo, the posterior GB elongated ∼ 2.5-fold and developed a neck region ∼ 5-fold narrower than the initial width (Figs. 2C, D). By contrast, the anterior GB remained on the ventral surface, narrowed only ∼ 40% and elongated by a similar factor. These contraction factors are consistent with experiment (32, 49, 55).

### The anterior GB is fluidized and reshaped by planar-polarized junctional myosin

In the following sections we examine the distinct mechanisms in the anterior and posterior GB, Fig. 2C. We define the posterior GB as the region where junctional myosin is small, i.e. the 40% most posterior cells at the onset of GBE (29, 66, 67) (Fig. 1C). The anterior contains the 60% most anterior cells.

Beginning with the anterior GB, in simulations this region underwent approximately incompressible convergent-extension with little deviation from rectangularity, Figs. 3A, D, F. While the anterior GB deformed as a whole, its cells remained undeformed: over 30 min, in simulations the anterior GB aspect ratio (AP:VL) increased ∼ 150% while cell aspect ratio decreased < 10%, similar to the experimentally reported cell aspect ratio decrease (36), Fig. 3B. This is the hallmark of fluidity, with cells acting like fluid particles that continuously reorganize spatially without individual deformation.

**Figure 3.**
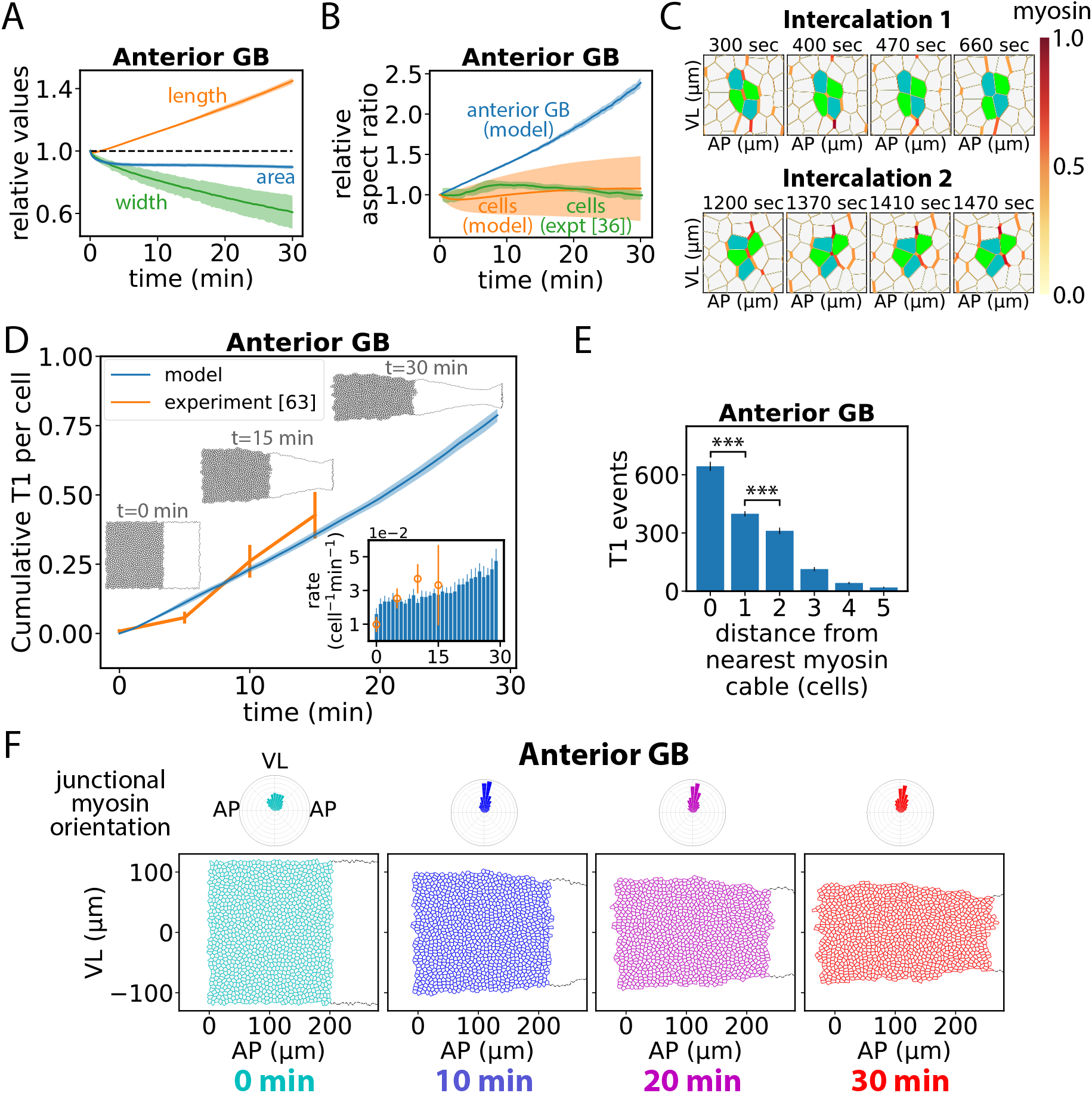
The anterior GB is fluidized and reshaped by planar-polarized junctional myosin. (A) Predicted relative anterior GB length (AP direction), width (VL direction) and area versus time. (B) Relative anterior GB and anterior cell aspect ratios (AP/VL) versus time. Experimental data replotted from ref. (36). (C) Examples of simulated T1 transitions in the anterior GB. The contracting (expanding) junction eliminated (spawned) by a T1 transition tends to have high (low) myosin. (D) Total number of T1 transitions per cell versus time in the anterior GB as predicted by the model (*n* = 10 emb**r**yos) and measured experimentally (replotted from ref. (63), *n* = 3 − 8 emb**r**yos). Inset: Data replotted as T1 rates per cell (bars: model; orange circles: experiment). Error bars, s.d. (E) Mean number of T1 transitions per junction in the anterior GB during 30 min of GBE versus distance from nearest myosin cable. Bars, s.d. (*n* = 10 emb**r**yos). (F) Evolution of anterior GB during GBE. Myosin orientational distributions are planar polarized. Shaded regions in (A), (B), s.d. (simulations, *n* = 10 embryos, ∼1127 anterior cells per embryo) or SEM (experiments, *n* = 3 emb**r**yos).

This fluid-like deformation was facilitated by cell intercalations, driven by junctional myosin in cables, Fig. 3C. Before myosin ramp-up the simulated GB is a solid (34) (see Model section). Following myosin activation, there was a brief fluidization period of ∼ 2.5 min during which the rate of cell intercalation by T1 processes increased from zero and reached steady state, Fig. 3D. The fluidization time is of order the duration of a single T1 event (76, 77). Thereafter, the intercalation rate remained roughly constant for ∼ 20 min, increasing gradually during the final ∼ 10 min. The predicted time dependence of intercalations reproduced experiment (63) during the first 15 min, Fig. 3D. Over 30 min, the ∼ 1130 anterior GB cells participated in 0.79 ± 0.02 intercalation events per cell (*n* = 10 simulations).

The driver of the convergent-extension flow was the planar polarity of the junctional myosin distribution which biased intercalation events: as VL-oriented junctions tended to have high myosin, cell intercalations typically involved loss of a VL-oriented junction and birth of an AP-oriented junction, so cells converged ventrally and diverged in the AP direction (Fig. 3C).

T1 intercalations occurred most frequently at myosin cable junctions (Fig. 3E), as expected since these junctions have greatest tension. This bias is seen experimentally (21, 34). In simulations lacking cable organization, where myosin was planar polarized but otherwise randomly distributed, the overall intercalation rates were virtually unchanged, but the GB elongation rate was ∼ 10% lower. The elongation was slower because neighboring T1 events often interfered laterally with one another (Fig. S4), whereas in cables all T1 events were laterally well separated. Given the smallness of this effect, we suggest cable organization serves primarily to define parasegment boundaries (see below).

In summary, planar polarized junctional myosin drives anisotropic fluidization of the anterior GB. To a good approximation, the anterior GB undergoes incompressible convergent-extension fluid flow.

### Anterior GB fluidization requires spatiotemporal myosin fluctuations

Anisotropic fluidization of the anterior GB driven by planar polarized junctional myosin was strongly enhanced by fluctuations. In accord with experimental data (21, 28), our model incorporates 30% junction-to-junction myosin fluctuations, with a 200 sec memory time. In simulations where these fluctuations were abolished, T1 transitions were ∼ 2-fold reduced, Fig. 4A. Bigger fluctuations or longer fluctuation memory time increased T1 transition rates and decreased cell aspect ratios, Figs. 4B, C. A similar trend was seen in a previous vertex model study in which increasing junctional tension fluctuations promoted fluidity (47).

**Figure 4.**
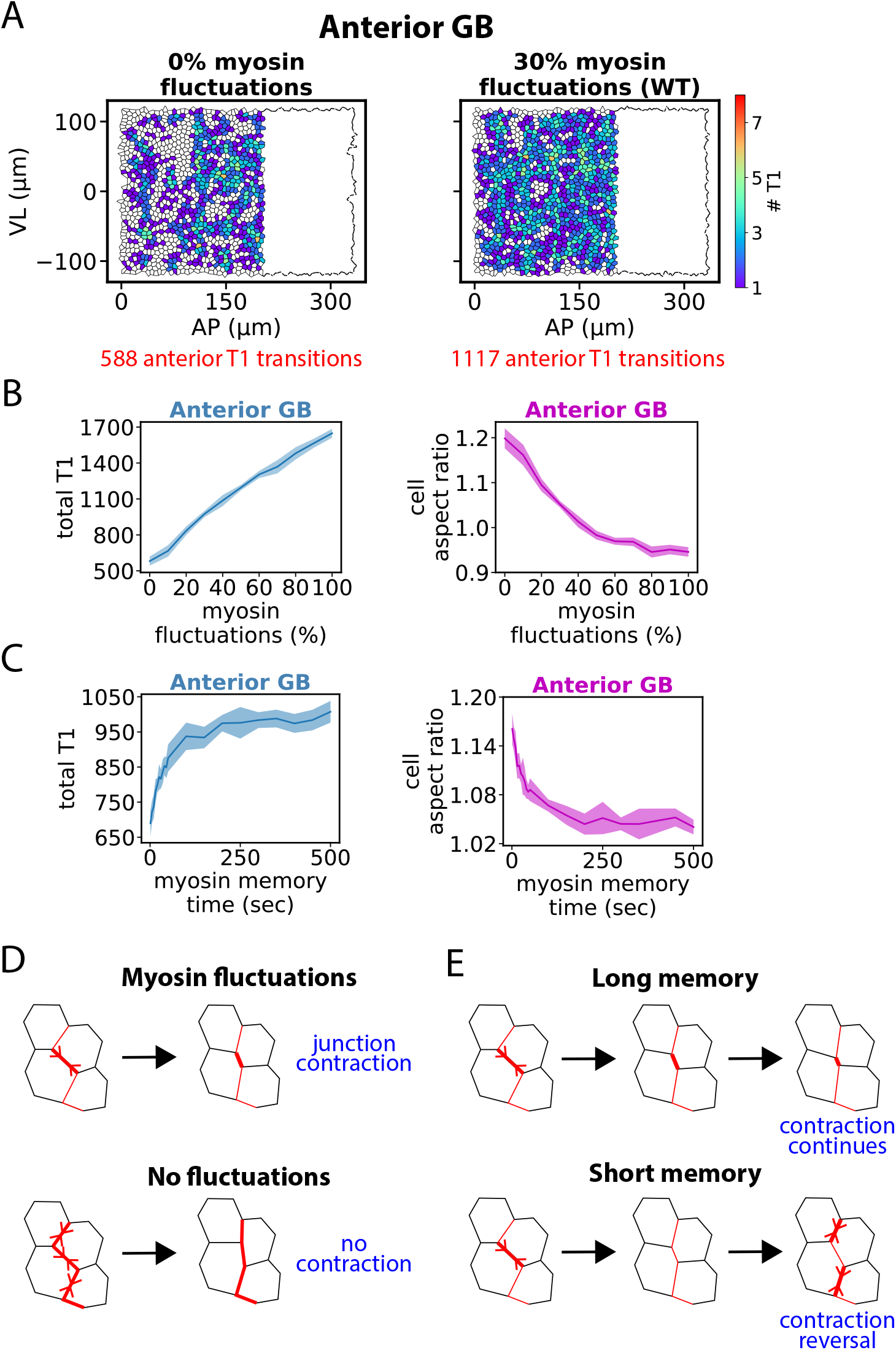
Anterior GB fluidization requires spatiotemporal myosin fluctuations. (A) Distribution of T1 transitions in the anterior GB during one simulation of the first 30 min of GBE. Transitions are mapped to cell locations at GBE onset. Abolishing temporal myosin fluctuations strongly lowers the T1 frequency (left). (B and C) Number of T1 transitions and mean cell aspect ratio (AP/VL) in the anterior GB during 30 min of GBE versus relative magnitude (B) or memory time (C) of myosin fluctuations (*n* = 10 embryos, ∼ 1127 anterior cells per embryo). Shaded regions, s.d. (T1 transitions) or SEM (aspect ratios). (D and E) Myosin fluctuations promote cell intercalations by generating tension gradients along myosin cables (D) provided the fluctuations are sufficiently long-lived (E).

Thus, fluidization is enhanced by long-lived fluctuations. This originates in the basic cell intercalation event, when a VL-oriented cell-cell junction contracts completely. This requires a stress gradient: the contracting junction must have higher myosin than the two similarly oriented neighboring junctions. Bigger fluctuations make this more likely to occur, Fig. 4D. Further, the gradient must persist sufficiently, lest the gradient is reversed before junction contraction is complete, Fig. 4E. Small or very transient fluctuations suppress fluidization by lowering the rate of intercalations that dissipate elastic energy, promoting elastic cell distortion and locking in elastic energy.

Thus, normal GBE requires large spatiotemporal fluctuations over and above ramp up. Ramping up junctional myosin does not in itself fluidize confluent tissue. Indeed, vertex modeling studies (71) and experiment (78) show that increasing junctional tension has the opposite effect, stabilizing the solid lattice structure.

### The posterior GB is elastically elongated by external forces whose range is friction-limited

Next we studied the evolution of the posterior GB. Unlike the anterior, little junctional myosin is present in the posterior (67) (Fig. 1C), suggesting its reshaping may be driven by external forces from neighboring PMG primordium tissue, Fig. 2A.

In simulations, GBE radically remodeled the posterior GB into a long, thin domain with a neck region, suited to wrapping around the embryo posterior (Figs. 5D, 2C, D). Deformation of the posterior GB was approximately incompressible, as for the anterior (Figs. 5A, 3A). However, its relative elongation was ∼ 3-fold greater (compare Figs. 5B, 3B), and compared to anterior cells posterior cells became much more stretched in the AP direction, with a mean aspect ratio of ∼ 2 (Figs. 5C, 3B). This large cellular deformation represents an elastic tissue deformation mode and is in quantitative agreement with experiment, Fig. 5C (30).

**Figure 5.**
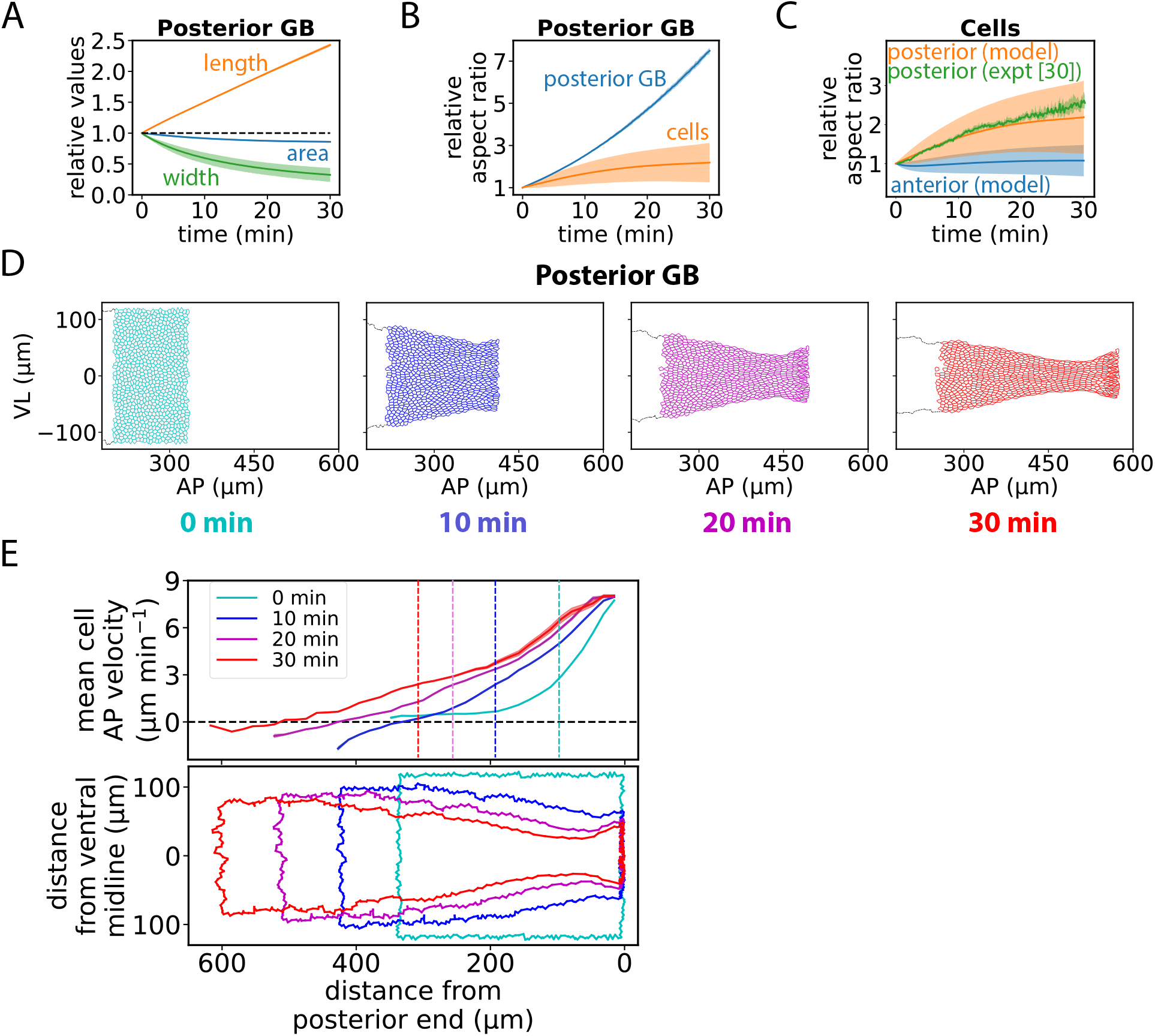
The posterior GB is elastically elongated by external forces whose range is friction-limited. (A). Relative posterior GB length (AP direction), width (VL direction) and area versus time. (B) The Posterior GB region and posterior GB cells undergo large deformations. Relative aspect ratios (AP/VL) versus time. (C) Posterior GB cells are much more strongly stretched in the AP direction than are anterior GB cells. Relative cell aspect ratios versus time. Experimental data replotted from ref. (30). (D) Evolution of posterior GB during a typical GBE simulation. (E) Due to frictional dissipation the mean cell velocity in the AP direction decays with distance from the posterior GB boundary with a penetration length that increases with time. Vertical dashed lines locate anterior/posterior boundary. The evolving GB shape is shown for reference (bottom). Shaded regions, s.d. (simulations in (A), (B), (C)) or SEM (experiment in (C)). Simulations in (A), (B), (C) used *n* = 10 embryos, ∼ 1127 anterior cells, ∼ 726 posterior cells per embryo.

Thus, the posterior and anterior GB undergo very different reshaping evolutions. The posterior GB is reshaped primarily by external forces acting on its posterior edge. But why do these forces have little effect on the anterior GB? In our model, this is due to frictional forces, likely from the vitelline membrane, that oppose cell motion. The posterior edge is pulled at ∼ 8 μm min^−1^ in the AP direction and pulls adjacent cells in the same direction but with reduced velocity due to frictional resistance (Fig. 5E). The AP velocity profile penetrates ∼ *n*(*t*) ≈ (*A*_0_*K*_*a*_*t*/*λ*)^1/2^ cells into the GB from the edge after time *t* (see Supplementary Information), a penetration depth of ∼ 30 cells after 20 min, about the posterior portion of the GB at that stage. Thus, the posterior pulling force penetrates the anterior GB only in the final ∼ 10 min of the 30 min of fast GBE, explaining the increased cell intercalation rates in the anterior GB during this period (Fig. 3D inset).

Overall, the posterior GB suffers severe elastic deformation due to the large external pulling force at its boundary, while friction screens the anterior GB allowing it to undergo mild myosin cable-mediated fluidization.

### Elastic stresses crystallize the posterior GB and activate cell intercalations that mediate remodeling

After thirty minutes of GBE, the posterior GB had a completely different shape to the anterior GB (compare Figs. 3F, 5D). The organization of cells also differed radically: while the anterior organization was essentially unchanged from the initial organization, the posterior GB acquired crystal-like order, with elongated cells arranged in VL-oriented stacks, Fig. 6A.

**Figure 6.**
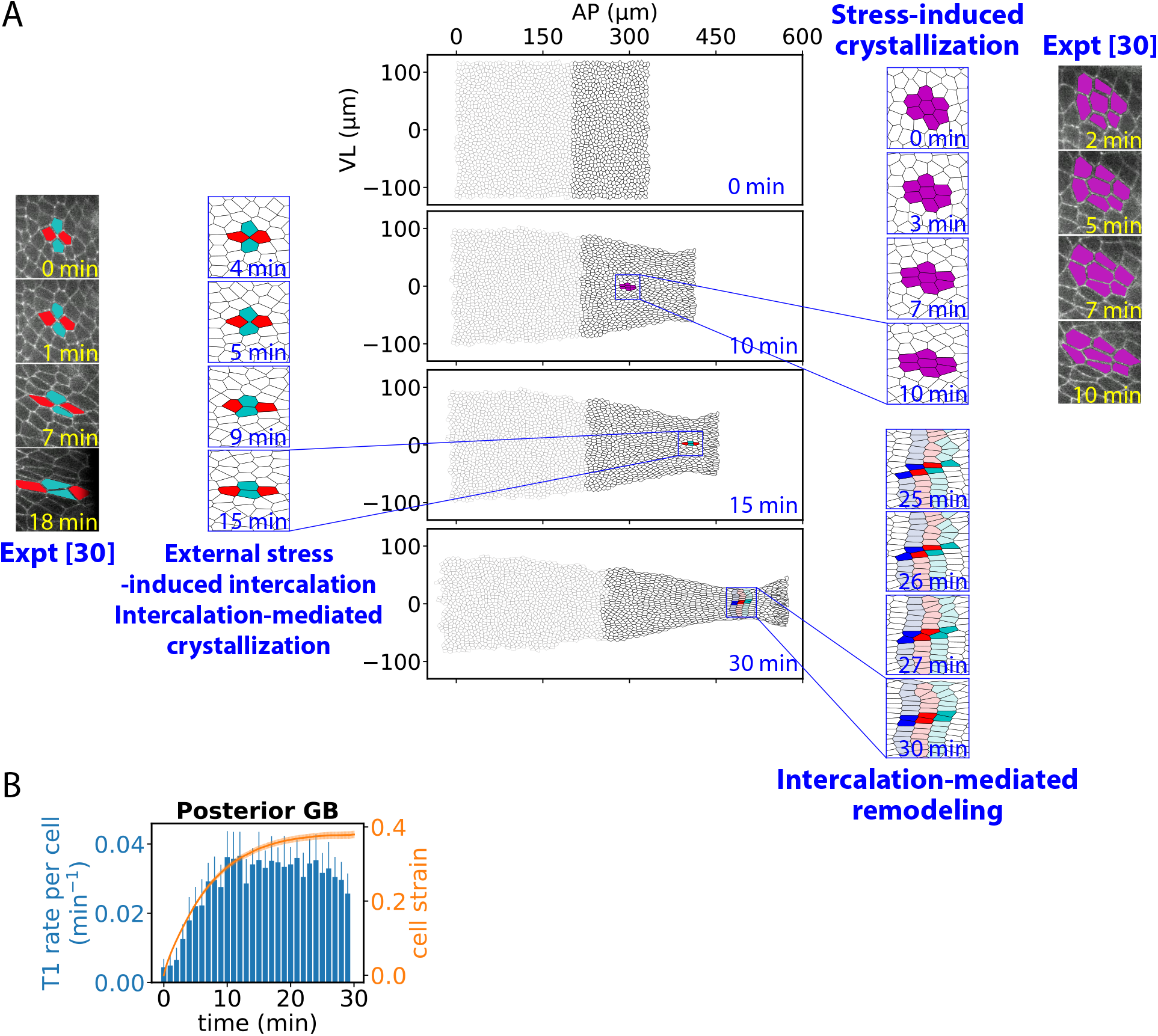
External forces activate cell intercalations that crystallize and remodel the posterior GB. (A) Evolution of GB during 30 min of GBE. Three key processes crystallize and reshape the posterior GB. External pulling forces stretch cells, directly inducing crystallization (stress-induced crystallization). Further stretching thins cells sufficiently to induce T1 cell intercalations (external stress-induced intercalation) that mediate further crystallization (intercalation-mediated crystallization). Stress-induced intercalations anneal crystalline defects and mediate plastic flow (intercalation-mediated remodeling). In the highlighted example, synchronized intercalations mediate shearing of cell stack assemblies (25 – 30 min). Microscopy images adapted from Ref. (30) show stress-induced and intercalation-mediated crystallization in embryos whose posterior GB was mechanically separated from the anterior (cadherin labelled). (B) T1 intercalation rates per cell and cell strains in the posterior GB versus time. Bars and shaded region, s.d. (*n* = 10 embryos, ∼ 726 posterior cells per embryo).

How did posterior crystallization come about? We observed two stages, Fig. 6A. (1) Stress induced crystallization. Due to the posterior pulling force, cells were stretched in the AP direction and became thinner in the VL direction due to their near incompressibility. The elongated cells tended to pack into ordered stacks, similar to rodlike molecules in a smectic (layered) liquid crystal phase (79). (2) Stress-induced intercalations that mediated crystallization. Continued narrowing in the VL direction contracted VL-oriented junctions, triggering T1 intercalations which further crystallized cells into stacks. T1-mediated stacking was efficient because cells had high tension from stage (1), so when a T1 event generated a new AP junction separating two cells, the junction rapidly elongated and the cells became bound along their new common junction.

As crystallization progressed, in response to the external pulling stresses the posterior GB continued to remodel due to a third key process, Fig. 6A. (3) T1-mediated remodeling. Remodeling was facilitated by T1-mediated stack reorganization, when stacks severed and reconnected to neighboring stacks. This mechanism annealed defects such as the vertex in Y-shaped stack configurations. Plastic flow of the solid was facilitated by synchronized T1-mediated restacking events that simultaneously reconnected neighboring stacks, effectively sliding stacks over one another in the AP direction (Fig. 6A, right).

Such crystallization processes in the posterior GB have been experimentally observed during GBE. In ref. (30), a laser cauterization procedure attached GB cells to the vitelline membrane along a line ∼120 μm from the posterior end of the embryo, positioning the line within the posterior GB and ∼ 10 μm from the anterior/posterior boundary (Fig. 2C). This mechanically isolated the two regions, ensuring the posterior GB was subject only to external boundary stresses from PMG invagination. In the posterior GB, cells were strongly stretched by the external stresses, leading to stress-induced crystallization (Fig. 6A, right) and stress-induced T1 intercalations that mediated crystallization (Fig. 6A, left).

In parallel with these remodeling events, the T1 intercalation rate in the posterior GB steadily increased with time as more cells transited to stages (2), (3) and participated in intercalations, Fig. 6B. T1-mediated stacking onset later for cells further from the boundary which had to wait longer for the domain of influence of the boundary force to penetrate to their location. As a result, after ∼ 15-20 min, the penetration time for the entire posterior GB, the T1 rate reached steady state. This contrasts with the anterior GB where intercalations enabled cells to continuously rearrange as a fluid with little cell shape change (Figs. 3B, C), and a steady state intercalation rate was reached much more rapidly (inset, Fig. 3D).

In summary, external stresses crystallize the posterior GB and activate cell intercalations that mediate defect annealing and flow of the solid-like tissue.

### Anterior fluid-like and posterior solid-like remodeling cooperatively reshape the GB

Both regions of the GB undergo convergent-extension, but the processes differ fundamentally. The anterior and posterior GB remodel as a fluid and a solid, respectively, yielding very different shapes. Working together, the two-state process reshapes the roughly rectangular GB at gastrulation onset into an asymmetrical tapering form by the end of the fast phase of GBE, with a wide anterior connecting to a much narrower elongated posterior that wraps around the embryo posterior (Figs. 2D, 6A, S5).

The posterior GB has much higher cell strains (Fig. 7A), as seen experimentally (32), and much higher elastic energies (Figs. 7B, S6). The degree of cell elongation and alignment during GBE can be quantified (33) by a shape parameter *Q* (Supplementary Information), which is always small in the anterior but in the posterior becomes large soon after GBE onset, reflecting cell alignment into local crystal-like order, Fig. 7D. Vertex models exhibit fluidization (loss of rigidity) above a critical value 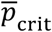 of the dimensionless shape parameter 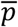 averaged over cells (see model section), but for a given shape tissue is more solid-like when cells are more aligned, so at higher *Q* the fluidization transition shifts (33) to higher 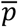. A theoretical expression (80) adapted to GBE experimental data (33) yielded the relation 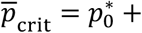 1.7 *Q*^2^ with 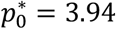. Applied to our simulations, this relation suggests the anterior GB rapidly fluidizes from the initial solid-like state, whereas the posterior GB is always solid-like due to large *Q* values (Fig. 7D). The distinct states are also reflected in cell streamlines, which in both regions are hyperbolic due to incompressibility (Supplementary Information and Fig. S8) but the anterior fluid-like trajectories have greater fluctuations (Fig. 7C).

**Figure 7.**
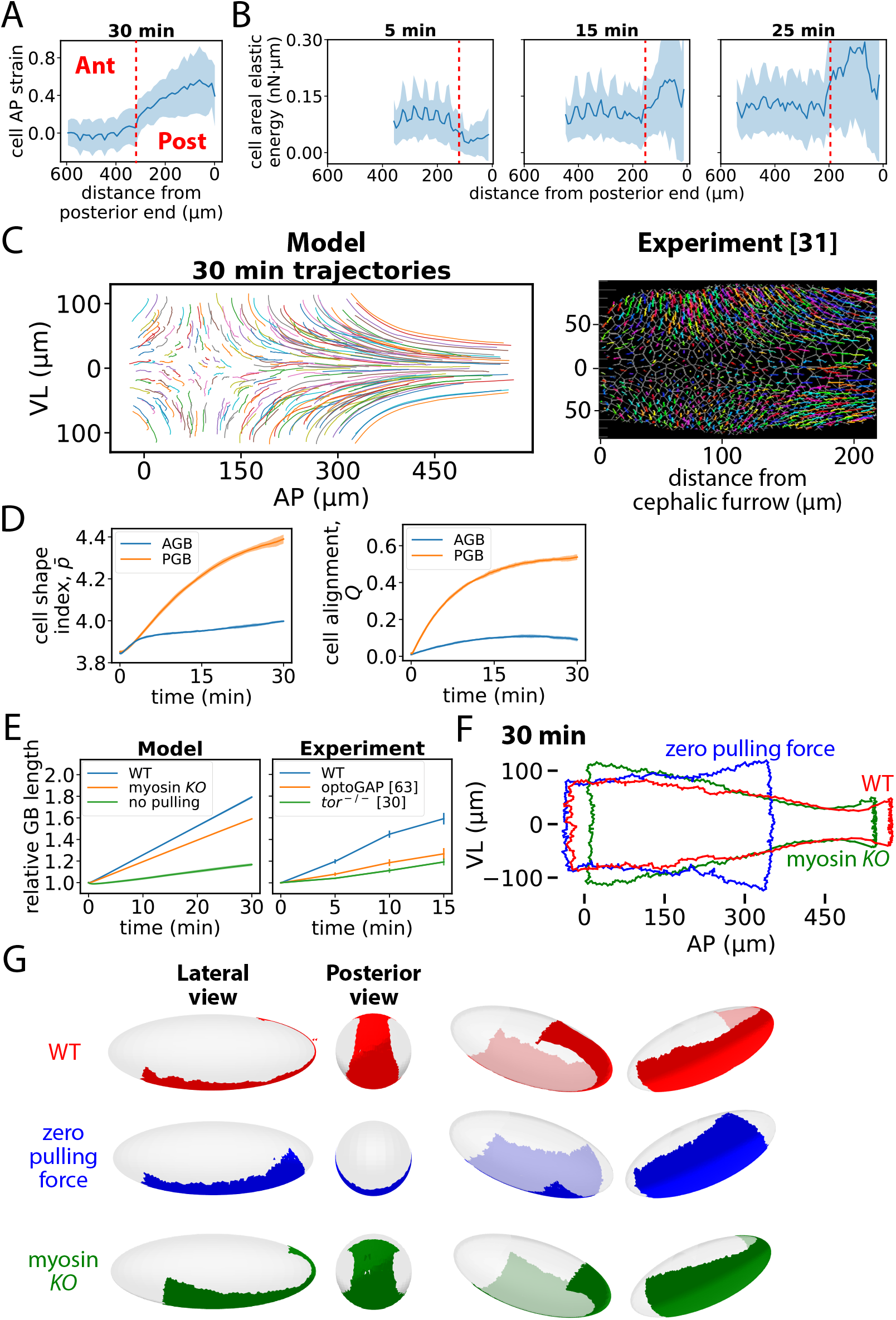
Anterior fluid-like and posterior solid-like remodeling cooperatively reshape the GB. (A) Profile of cell strains in the AP direction. (B) Profiles of cell areal elastic energies. (C) Selected simulated cell trajectories (left). Experimental cell trajectories (4 min duration, starting 10 min after GBE onset) replotted from ref. (31) (right). (D) Mean cell shape index, *p* (left) and cell alignment, *Q* (right) versus time in simulations (*n* = 10 emb**r**yos; shaded regions, s.d.). (E) Left: relative GB length (AP direction) versus time in wild-type, myosin knockout and zero posterior pulling force simulations (*n* = 10 embryos; curve thicknesses, s.d.). Right: Experimental data for the same quantities in wild-type embryos, optoGAP embryos and embryos from torso-mutant females, replotted from refs. (30, 63) (*n* = 3 − 8 embryos; bars, SEM). (F) GB shape after a wild-type, a myosin knockout and a zero posterior pulling force GBE simulation. (G) GBs of (E) visualized on embryo surface. In (A) and (B) red dashed lines indicate anterior/posterior boundary, shaded regions denote s.d. and *n* = 10 embryos of 1853 cells were simulated.

The complimentary roles for anterior and posterior remodeling are highlighted by simulated and experimental mutant phenotypes. In simulations with no myosin, GB elongation was ∼ 25% reduced, compared to a ∼ 50% reduction in experiments where myosin activation was suppressed using optoGAP, Fig. 7E. In these experiments, expression of a photosensitive RhoGAP reduced GB elongation ∼15% even without light, suggesting the true reduction due to optical inactivation is closer to the simulated value. Without myosin, the simulated GB still has its extended posterior but little convergent-extension of the anterior occurs, Figs 7E, F. In simulations with zero posterior pulling force, elongation was ∼ 75% lower, similar to the ∼ 60% reduction in experiments with posterior midgut invagination abolished (30, 63), Fig. 7E. Lacking an elongated posterior, the GB no longer extends to the dorsal surface, Figs. 7F, G.

### Myosin cables maintain parasegment organization by selective fluidization of the boundaries

We found that the VL-oriented cable organization (21, 23) of junctional myosin in the anterior GB had relatively little effect on the rate of convergent-extension (Figs. 1D, 3E, S4), suggesting some other function. Indeed, during the fast (21) and slow (81) phases of GBE the cables were proposed to maintain the organization of parasegments, fundamental developmental units crucial to body plan organization (82). In ref. (21) it was shown that cells do not cross the boundaries during the fast phase, and it was argued that cell mixing is prevented by the high line tension of the myosin cables, which lie at the parasegment boundaries. However, the underlying mechanisms have not been elucidated.

In the present model, following experiment (21) myosin cables are initially positioned periodically along the GB, with one strong cable every five cell rows, defining the initial parasegment configuration (Fig. 8A). Over 30 min of GBE this organization was remarkably well preserved: no cell transmissions occurred, reflected by a mean parasegment interface roughness of ∼ 0.3 cells and an absence of cell islands displaced from their native parasegments (Fig. 8A). The mechanism is that most T1 transitions in the anterior GB are driven by myosin cables, so they occur at parasegment boundaries (Figs. 3E, 8C): a cable junction contracts to zero, leaving the cable intact while pushing the cells away from the boundary, guaranteeing they remain in their home parasegments (Fig. 8B, red cells).

**Figure 8.**
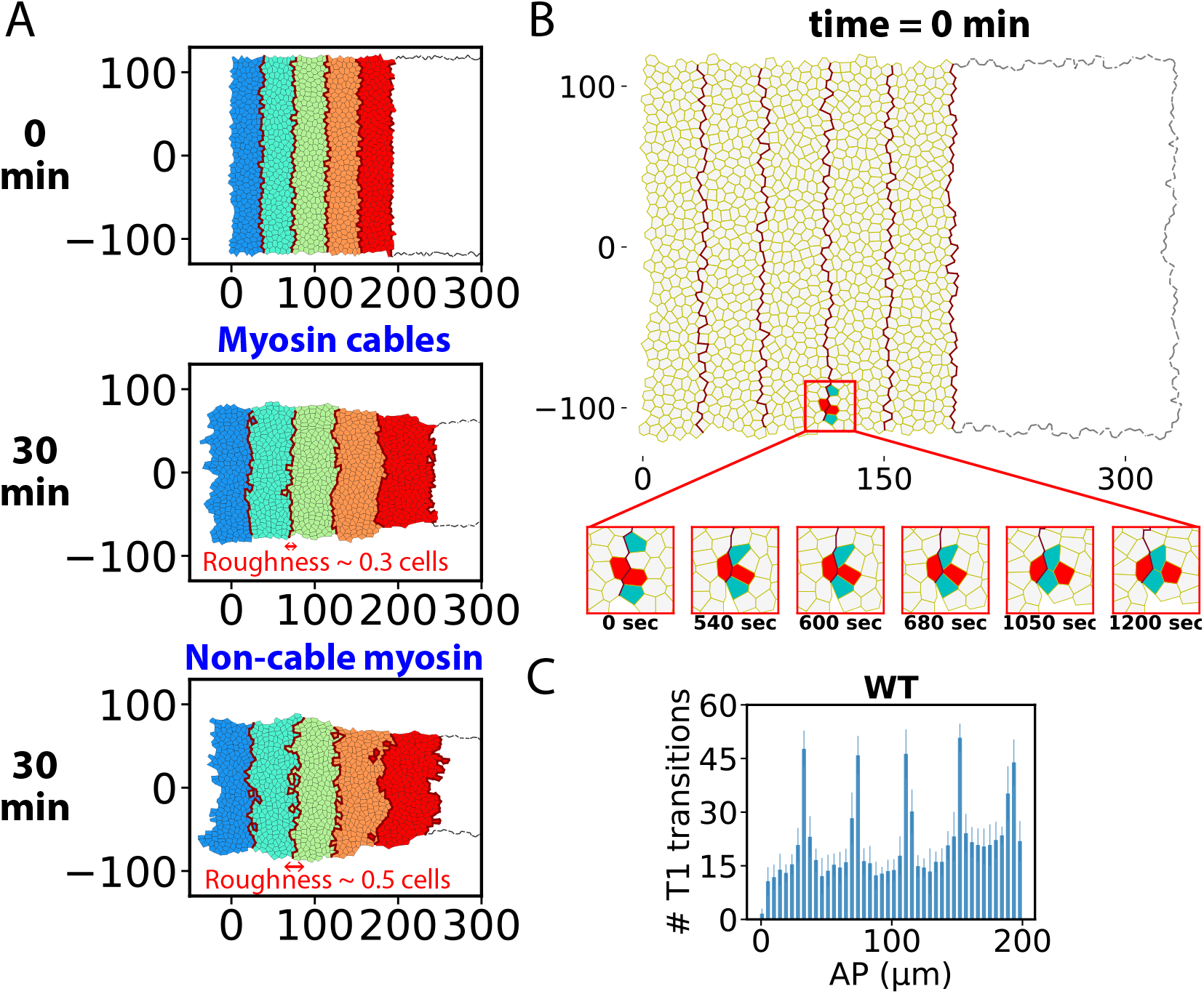
Myosin cables confine cells to their home parasegments by selectively fluidizing parasegment boundaries. (A) Evolution of parasegments in the anterior GB after 30 min of GBE simulation with normal myosin organization (cables, wild-type) or with random non-cable myosin organization. Without cables cells migrate from their home parasegments and parasegment boundaries are rougher. (B) A typical T1 transition at a myosin cable that coincides with a parasegment boundary in the anterior GB during a wild type GBE simulation. The transition preserves cable integrity and maintains parasegment organization by pushing lateral cells into their home parasegments (red cells). For clarity, only the anterior GB is shown. (C) Spatial distribution of numbers of T1 transitions following a 30 min wild-type GBE simulation (*n* = 10 emb**r**yos, bars are s.d.) The five peaks correspond to five parasegment boundaries. Position coordinate was rescaled to initial GB.

In simulations with random planar polarized but non-cable myosin organization, T1 transitions often occurred away from parasegment boundaries so the protective cable effect was lost. Numerous cells crossed the boundaries which became badly distorted (Fig. 8A, bottom right). For the non-cable run of Fig. 8A, 19 ± 6.2 cell islands (10 embryos) crossed into neighboring parasegments and the interface roughness increased to ∼ 0.5 cells.

Thus, organizing myosin into elongated cables at parasegment interfaces localizes T1 transitions and fluidization to the interfaces. Selective fluidization of the interfaces allows parasegments to slide relative to one another with little interfacial disruption, so parasegment identity is maintained.

## Discussion

### Germ-band extension: a two-state tissue reshaping strategy

It is known experimentally that cell shapes, intercalation rates and junctional myosin levels during GBE are very different in the anterior and posterior GB (see (21, 29-32, 34) and Figs. 1, 3, 5, 6). Using a vertex model framework, we concluded that these distinct properties correspond to distinct tissue states in the two regions: a slowly deforming anterior fluid, and a posterior solid undergoing ∼ 2-fold faster elastic elongation and narrowing (Fig. 2). Thus, while *Drosophila* GBE is a classic instance of convergent-extension, it consists of two qualitatively distinct convergent extension processes that occur in parallel, with different extension rates and entirely different cell intercalation mechanisms.

Simultaneous activation of the two states allows GBE to accomplish the challenging task of remodeling the roughly rectangular GB into a bottle shape, requiring moderate anterior extension and a severely extended posterior neck that wraps around the embryo posterior onto the dorsal surface. The crystal-like ordering of cells in the solid-like posterior tissue may confer the advantage of a highly regular boundary with the laterally adjacent amnioserosa tissue, as boundaries in model simulations were far more irregular in the fluid anterior (cf. Figs. 3F, 5D). In principle, the high posterior strain rate could be accomplished by a strongly driven posterior fluid, but we suggest the high junctional myosin loading required might excite instabilities (related to boundary irregularity) that destabilize the thin, extended posterior shape. Further, cell intercalations may be difficult to activate at the high curvature posterior end where cells must adopt wedge shapes, inhibiting junction contraction on the expanded apical surfaces.

### Internal fluctuating junctional stresses fluidize tissue

From solid state, tissue can be fluidized by random cell intercalations driven by spatiotemporally and orientationally stochastic myosin fluctuations at cell-cell adherens junctions (83). Any given junction may then randomly close down at any instant due to a fluctuation that boosts its myosin sufficiently to overcome the restraining effect of neighboring junctions, giving an intercalation, Fig. 4D (47). In *Drosophila*, this situation is realized in the pupal notum where fluidization is thought to endow tissue with robustness in the face of cell divisions and other disruptions (78, 84). Cells are then, in effect, randomly and isotropically displaced by intercalations they participate in. Provided the intercalations are spatially dilute the stochastic cell trajectories are expected to be independent, similar to those of molecules in a fluid.

GBE uses this mechanism to liquefy the anterior GB, with the additional feature that the junctional myosin is anisotropically distributed. The fluidity of the anterior is manifested by the fact that a steady state is reached, in contrast to the solid-like posterior: following an initial ∼ 2 min fluidization episode when junctional myosin ramps up and intercalation rates increase from zero, the intercalation rate becomes roughly constant (35, 63) (Fig. 3D inset). Unlike the pupal notum, the junctional myosin is orientationally biased in VL-oriented cables (21, 23) (Figs. 1D, 3C) which bias the orientation of T1 intercalations so cells converge (diverge) in the VL (AP) directions (34), producing convergent-extension (Figs. 3C). At any instant ∼ 30 intercalations are occurring in the ∼ 1100 cell anterior GB, implying that intercalations are separated by ∼ 10 cells, sufficiently dilute that cells can recover their shape. Indeed, experimentally and in model simulations anterior cell shape is little altered from rest (Figs. 3B, F).

Two of our key conclusions are as follows. (i) Junctional myosin fluctuations are required for high anterior fluidity. Planar polarity alone is insufficient, as junction rotation during intercalations tends to block completion (Fig. 4D). Without myosin fluctuations, intercalations were ∼ 2-fold lowered and cell shape was strongly perturbed (Fig. 4B). (ii) Junctional myosin planar polarity plus uncorrelated myosin fluctuations are sufficient for anterior fluidization and convergent extension, without need for intercellular correlations. Planar polarity ensured that a new junction spawned by an intercalation is very likely to grow, since such junctions have strongly AP-biased orientation and thus much lower junctional myosin than the VL-oriented junctions whose closing down initiated the intercalation (Fig. 3C). Thus the intercalation is very likely to complete, without need for correlations in junction properties of lateral cells (84).

Our model neglected junctional adhesion planar polarity (26, 27), but we expect our qualitative conclusions to be unaffected since experimentally E-cadherin levels were found to have little effect on intercalation rates or net GB elongation (85). Implicitly we assume fast turnover of adhesion components so junction dynamics are controlled by junctional myosin, consistent with junction shortening times of ∼ 5 min (76) compared to reported cadherin turnover timescales of ∼ 2 min or significantly less due to mechanosensitivity (86, 87).

### External stress drives cell intercalation, tissue crystallization and plastic flow in the posterior GB

The posterior GB is reshaped by external stresses from adjacent tissue acting on the posterior boundary whose effect is very different to that of internal fluctuating stresses from junctional myosin cables, which are absent from the posterior. Consistent with experiment (30), we found external stresses drive crystallization and plastic flow of the posterior GB tissue mediated by cell intercalations whose origin is quite different to that of anterior junctional stress-driven intercalations.

In the first 5-10 min of GBE, in model simulations the posterior GB tissue behaved similarly to an elastic solid, the external forces stretching individual cells and the tissue as a whole in the AP direction (Fig. 5), as seen experimentally (30-32). As cells resist volume change (60), AP stretching generates high stresses tending to thin cells in the VL direction. These stresses drive local crystal-like ordering by aligning the elongated cells, allowing cells to thin and relieving the stresses and associated elastic energy (stress-induced crystallization, Fig. 6A). This is seen experimentally in the posterior GB (30).

Within ∼ 5-10 min, elastic stress-driven cell thinning due to continued GB elongation is sufficient to close down VL-oriented junctions and activate cell intercalations (external stress-induced intercalations, Fig. 6A). A new AP-oriented junction created by such an intercalation rapidly and irreversibly expands in the AP direction, relieving elastic stress and energy and providing a new interface for two cells to align and further advance the crystal-like tissue ordering, (intercalation-mediated crystallization, Fig. 6A). As crystallization progresses, stress-induced intercalations cooperatively anneal defects by collective rearrangements such as sliding of crystalline stacks, analogous to defect annealing by crystal plane sliding in inanimate crystalline materials (88, 89) (intercalation-mediated remodeling, Fig. 6A).

Thus, the posterior GB undergoes convergent extension deformation despite its solid and progressively crystalline character. Initially, this is primarily simple incompressible elastic solid deformation. After 5-10 min, the crystal-like tissue undergoes plastic flow mediated by stress-induced intercalations. Such intercalation-mediated crystallization and remodeling processes have been observed experimentally (30). The initially slow increase in posterior intercalation rates (Fig. 6B) reflects the time lag before cells are sufficiently stretched and narrowed to activate intercalations, in contrast to their almost immediate onset in the anterior (Fig. 3D) due to internal junctional myosin. Another difference is that while anterior cell shape is virtually unaltered, after 30 min of GBE posterior cells are highly elongated, with aspect ratio ∼ 2 (Fig. 5C). Significantly, this is much less than the ∼ 7 aspect ratio of the posterior GB as a whole (Fig. 5B) due to plastic flow of the solid-like tissue.

### Wound healing is another two-state tissue reshaping process

Two-state GBE has remarkable parallels with wound healing (90, 91). Wounding in the *Drosophila* wing disc evokes a healing response with an initial ∼ 20 min fast phase when a contractile actomyosin cable at the inner wound margin exerts inward radial stresses. These are effectively external stresses as they act on the inner boundary of the tissue, analogous to the external stresses from PMG invagination acting on the posterior GB boundary. The leading-edge cells are radially elongated and azimuthally narrowed by these external stresses, provoking their radial alignment in an annular stack at the wound boundary, similar to stress-induced crystallization in the posterior GB, Fig. 6A. In the subsequent >200 min slow phase, continued external stress activates cell intercalations at the wound edge. The intercalations continuously decrease the number of leading-edge cells without disturbing their regular crystal-like annular organization, maintaining a regularly shaped wound edge for good wound closure. This process is similar to intercalation-mediated crystallization in the posterior GB, Fig. 6A.

Similar to the liquid-like anterior GB, evidence suggests the epithelium away from the wound edge is fluid-like. Tissue fluidity in this region is required for normal wound closure: with elevated junctional myosin, closure failed and laser ablation measurements suggested increased tensions, consistent with higher tissue rigidity impeding closure, while wound closure was faster with reduced myosin and tension (90). These results suggest tissue fluidity in the epithelium facilitates healing. Consistent with these experiments, in the same study vertex model simulations with lowered tension away from the wound edge showed increased tissue fluidity and faster wound closure.

Thus, wound healing is another process using a mixed fluid-solid strategy, in this case for the delicate tissue reshaping challenge of healing in a circular geometry rather than linear as in GBE. The inner solid-like wound edge and outer fluid-like epithelium play similar roles to the posterior and anterior GB, respectively.

## Acknowledgements

This work was supported by the National Institute of General Medical Sciences of the National Institutes of Health under Award Number R01GM086731, and by the Human Frontier Science Program (Grant No. RGP001/2023). The content is solely the responsibility of the authors and does not necessarily represent the official views of the National Institutes of Health.

## Author contributions

B.O’S. conceived the study. B.O’S., T.Z. and H.Z. designed the model and performed the mathematical modeling. B.O’S. and T.Z. analyzed the data. B.O’S., T.Z. and H.Z. wrote the manuscript.

## Data and Code Availability

Simulation results supporting the findings of this paper, and codes to perform the simulations, to analyze the data, and to generate the technical figures are available in the GitHub repository [https://github.com/OShaughnessyGroup-Columbia-University/germ-band_extension.git].

