## Supplementary Material for "*Drosophila* germ band extension: a two-state reshaping mechanism"

**for**

**This PDF file includes:**

Supporting text

Figures S1 to S10

SI References

**Supporting text**

**Junctional actomyosin tensions**

Our simulation assumes that junctional actomyosin tension is proportional to the junctional myosin level. Therefore, junctional actomyosin tensions were estimated based on the myosin signals measured in wild-type embryos. The spatiotemporal tension profile $T_{i, k}^{\mathrm{myo}}$ for a cable junction is modeled as

$T_{i, k}^{\mathrm{myo}}=\left( 1+\epsilon_{i, k}\left( t \right) \right)\overline{T}_{i, k}^{\mathrm{myo}}$,

(S1)

where

$\overline{T}_{i, k}^{\mathrm{myo}}\boldsymbol{=}\overline{T}^{\mathrm{myo}}c_{ik}f_{ik}g_{ik}\cos\theta_{i, k}$,

(S2)

In Eq. (S1), $\epsilon_{i, k}\left( t \right)$ represents the temporal fluctuation relative to the mean tension, $\overline{T}_{i, k}^{\mathrm{myo}}$, at an individual junction $\left( i, k \right)$, which accounts for the location and the orientation of the junction. In Eq. (S2), $\overline{T}^{\mathrm{myo}}$ denotes the mean tension at a cable junction oriented parallel to the ventral-lateral (VL) direction (Table 1). $f_{ik}g_{ik}$ describes the spatial envelope of junctional myosin, with $f_{ik}$ and $g_{ik}$ being the mean values of the spatial envelopes $f\left( y_{n} \right)$ and $g\left( x_{n} \right)$ of the cells that junction $\left( i, k \right)$ belongs to, respectively. $f\left( y_{n} \right)$ and $g\left( x_{n} \right)$ depend on the position $\left( x_{n}, y_{n} \right)$ of the center of mass of cell $n$, with $x$ and $y$ denoting the anterior-posterior (AP) and the VL coordinates. Myosin planar polarity is captured by $\cos\theta_{i, k}$ term, reproducing the experimentally observed dependence of myosin intensity on junctional orientations(1).

**Myosin envelope**

Experimental data show that, in the VL direction, junctional myosin is enriched along the ventral midline and decays toward the lateral edges of the germ band (GB)(2) (Fig. S10). In the AP direction, the myosin level remains approximately constant over the anterior 60% of the GB and decays to nearly zero in the posterior GB(3) (Fig. 1C). To capture these spatial features, the VL envelope, $f\left( y_{n} \right)$, was extracted from experimental measurements of the global myosin level 5 min after GBE onset(2), while the AP profile, $g\left( x_{n} \right)$, was defined to reproduce the experimentally observed anterior–posterior decay. Myosin levels in the VL direction were rescaled by their maximum intensity, giving

$f\left( y_{n} \right)=e^{-1/2\left( 2y_{n}/L_{\mathrm{VL}}^{0} \right)^{2}}$,

(S3)

$g\left( x_{n} \right)=\left\{ \begin{aligned} 1, &x_{n}\leq aL_{\mathrm{AP}}^{0} \\ e^{-1/2\left[ \left( x_{n}-aL_{\mathrm{AP}}^{0} \right)/L_{\mathrm{AP}}^{0} \right]^{2}}, &x_{n}>aL_{\mathrm{AP}}^{0} \end{aligned} \right.$,

(S4)

where $L_{\mathrm{AP}}^{0}$ and $L_{\mathrm{VL}}^{0}$ are the initial AP and VL lengths of the GB, respectively, and $aL_{\mathrm{AP}}^{0}$ marks the boundary between the anterior and posterior GB, with $a=60\%$. This formulation ensures that $f\left( y_{n} \right)$ exhibits a bell-shaped profile in the VL direction, peaking at the ventral midline (Fig. S10), while $g\left( x_{n} \right)$ remaining flat in the anterior GB along the AP direction before dropping sharply at the anterior–posterior boundary.

**Myosin fluctuations**

Fluctuations in junctional myosin levels are a crucial aspect of our model whose magnitude is governed by the parameter $\epsilon_{i, k}\left( t \right)$, which represents the size of the fluctuations relative to the mean myosin value at individual junctions. Here we outline the procedure we employed to estimate the value of $\epsilon$ from experimental measurements. Using the reported experimental myosin profile from ref(1)., we first extracted the mean myosin level for the selected junction. We then assumed that for each individual junction $(i, k)$, the myosin level follows a normal distribution with a mean equal to the expected myosin level, $\overline{T}_{i, k}^{\mathrm{myo}}$, after applying the myosin spatial envelope, $f\left( y_{ik} \right)g\left( x_{ik} \right)$, myosin planar polarity, $\cos\theta_{i, k}$, and the cable effects. From the curve for the selected junction, we estimated the standard deviations of myosin levels relative to the mean. We assumed that the relative fluctuation amplitude $\epsilon$ (standard deviation divided by the mean) is the same for all junctions, which is approximately $30\%$. Thus, as a representative value for this experimental data, we used a value of $\sigma=30\%$ in our model.

**The stochastic factor** $\boldsymbol{\epsilon}$ **to represent myosin signals**

The time correlation of $\epsilon_{i, k}$ at a given junction $\left( i, k \right)$ has the form

$$\left\langle\epsilon_{i, k}\left( t' \right) \epsilon_{i, k}\left( t'+t \right) \right\rangle_{c}=\left\langle{\epsilon_{i, k}}^{2} \right\rangle_{c}\exp\left( -\frac{t}{\tau} \right).$$

In other words, $\epsilon_{i, k}(t)$ is the trajectory of a Brownian particle within a harmonic potential – the Ornstein-Uhlenbeck process. The amplitude of the fluctuations is taken to be $\left\langle{\epsilon_{i, k}}^{2} \right\rangle_{c}^{1/2}=\sigma=0.3$ for both wild-type and no-pulling simulations, estimated from experiment^1^. For both cases, we took the correlation timescale to be $\tau=200 s$ to match the experimentally reported fluctuations^1^.

**Posterior pulling velocity and force**

In simulations, the pulling from the PMG primordium as it invaginates concurrently with GBE is represented by a moving boundary condition with constant velocity, $v_{\mathrm{pull}}=8 \mu m \min^{-1}$, measured in wild-type embryos^2–4^. The force required for a vertex at the posterior end of the germ band to move at $v_{\mathrm{pull}}$ is calculated based on the force balance at the vertex:

$\mu\sum_{k}^{\left( i \right)} \frac{d\mathbf{L}_{i,k}}{dt}-\lambda\mathbf{v}_{i}+\frac{\partial H}{\partial\mathbf{r}_{i}}+\sum_{k}^{\left( i \right)} \mathbf{T}_{i,k}^{\mathrm{myo}}=-\boldsymbol{F}_{\mathrm{pull}, i}$,

(S6)

where the Hamiltonian *H* has the form of Eq. 2. Pulling forces are applied only to the vertices at the posterior boundary of the GB as they are directly connected to the PMG primordium. The total pulling force on a posterior cell is calculated as the sum of the forces applied to its associated vertices:

$\boldsymbol{F}_{\mathrm{pull}, n}=\sum_{i}^{(n)} k\boldsymbol{F}_{\mathrm{pull}, i}$,

(S7)

where $k=1$ if vertex $i$ is associated with cell $n$ only, or $k=1/2$ if vertex $i$ is shared with one neighboring cell besides $n$.

In simulations with no posterior pulling force, $v_{\mathrm{pull}}$ is set to zero.

**Mapping the 2D germ band to the surface of the 3D ellipsoid**

To approximate the 3D embryo geometry, we mapped the 2D GB sheet onto the surface of a prolate ellipsoid representing the *Drosophila* embryo (Fig. SX). The ellipsoid dimensions were estimated from experimental side-view images (with scale bar), yielding a total AP length of 490 µm and a maximal dorsoventral (DV) diameter of 180 µm. The ellipsoid is centered at the origin, with semi-axes $a_{X}=245 \mu m$, $a_{Y}=90 \mu m$, and $a_{Z}=90 \mu m$. The surface of the ellipsoid thus satisfies

$\frac{X^{2}}{{a_{X}}^{2}}+\frac{Y^{2}}{{a_{Y}}^{2}}+\frac{Z^{2}}{{a_{Z}}^{2}}=1$.

The positions of vertices in the 2D GB are given by coordinates $\left( x, y \right)$, corresponding respectively to the AP and VL directions in the tissue. Before GBE, the anterior end of the simulated GB is located at $x=0$. In the embryo, however, the anterior end lies at approximately one-quarter of the embryo length from the anterior pole. To align the two coordinate systems, we shifted the 2D GB coordinates by a constant offset $a=125 \mu m$:

$x'=x-a$.

The shifted coordinate $x'$ measures the arclength distance along the ventral midline of the ellipsoid, with $x'>0$ directed posteriorly and $x'<0$ anteriorly.

Each vertex in the 2D germ-band sheet was then projected onto the ellipsoidal surface following a two-step geodesic mapping.

1. **AP mapping**.

Starting from the ventral center $\left( X, Y, Z \right)=\left( 0, 0, -a_{Z} \right)$, the vertex “walks” along the ventral midline of the ellipsoid in steps of $0.01 \mu m$ along the local tangent direction, until the total arclength equals $\left| x' \right|$. This defines a midline point $\left( X_{x}, 0, Z_{x} \right)$.

2. **DV displacement on the ellipsoid surface**

At $\left( X_{x}, 0, Z_{x} \right)$, we first computed the surface normal vector $\mathbf{n}$, which defines the local tangent plane of the ellipsoid. Within this plane, a direction $\mathbf{v}_{y}$ perpendicular to the ventral midline tangent was chosen as the initial DV direction. The vertex was then iteratively displaced along this direction in $0.01 \mu m$ steps on the ellipsoidal surface.

After each displacement, the new local surface normal $\mathbf{n}_{\mathrm{new}}$ was recalculated. Because the surface is curved, the new displacement direction must be adjusted to remain tangent to the surface while preserving smooth continuation from the previous step. To achieve this, we first computed the unit normal $\mathbf{p}$ of the plane spanned by $\mathbf{n}_{\mathrm{new}}$ and the previous displacement direction $\mathbf{v}_{y}$:

$\mathbf{p=}\frac{\mathbf{n}_{\mathrm{new}}\boldsymbol{\times}\mathbf{v}_{y}}{\sin\left\langle\mathbf{n}_{\mathrm{new}}, \mathbf{v}_{y} \right\rangle}$.

The updated displacement direction was then redefined as the cross product between $\mathbf{n}_{\mathrm{new}}$ and $\mathbf{p}$:

${\mathbf{v}_{y}}^{'}=\mathbf{n}_{\mathrm{new}}\times\mathbf{p}$.

This ensures that the new direction ${\mathbf{v}_{y}}^{'}$ lies in the same plane as the previous step and remains orthogonal to the local surface normal, thereby constraining motion to the ellipsoid surface while maintaining a continuous orientation. The process was repeated until the cumulative surface displacement reached $\left| y \right|$, with the sign of $y$ determining whether the vertex moved toward positive or negative $Y$.

**Determining myosin cable positions and cell–cable distances**

In our simulation, myosin cables evolve over time and become increasingly irregular due to tissue deformation and cell shape changes during GBE. T1 transitions consume cable-associated junctions, causing those cable segments to disappear. To identify the positions of myosin cables at each time point, we selected all remaining cable junctions and classified them into 8 groups based on their AP positions, corresponding to the 8 initial myosin cables. For this classification, we applied the K-means clustering algorithm (from scikit-learn), assigning each junction a label indicating its cable membership. Junctions with the same label are considered as part of the same myosin cable. The AP position of each cable was then computed as the mean AP coordinate of its constituent junctions. A cell is considered a cable cell if the distance between its center of mass and the nearest cable is less than one average cell length.

**Both anterior and posterior GB cell trajectories are hyperbolic**

Experiments show that the GB tissue deformation approximately preserves area. From the experiments of ref. (4), the global strain rates in the AP and VL directions (x and y directions, respectively) vary over time, but remain approximately equal and opposite to one another, with a maximum difference of $0.04 \min^{-1}$, at GBE onset (Fig. S8A). For a rectangular GB of length $L$ ($x$direction) and width $W$ ($y$ direction) shortening in the $y$ direction and elongating in the $x$ direction, equal and opposite global strain rates $\dot{L}/L=-\dot{W}/W$ implies the area is constant in time. Thus, the GB undergoes approximately area-conserving convergent-extension, consistent with the ~ 10% net area decrease reported in ref.(5)

Similar behavior is observed in our simulations, with almost equal and opposite strain rates and conserved area (Fig. S8A). Assuming spatially uniform strain rates within the GB, the velocity $\boldsymbol{v}$ of a cell at $(x,y)$ would have components $v_{x}=\alpha\left( t \right)x$ and $v_{y}=-\beta\left( t \right)y$ where $(x,y)=(0,0)$ is the location of zero velocity. Incompressibility requires $\boldsymbol{\nabla}\cdot\mathbf{v}\boldsymbol{=}0$, so $\alpha\left( t \right)=\beta\left( t \right)$. Thus along the trajectory of a given cell, $d(xy)/dt =v_{x}y+{xv}_{y}= 0$, so cells would follow hyperbolic trajectories in which the product $x\left( t \right)y(t)$ has the initial value for that cell.

We measured cell trajectories to test if they are hyperbolic. Qualitatively, simulated and experimental trajectories are close to one another and have hyperbolic character (4, 6) (Figs. 7C, S8B). We grouped cells with similar initial coordinate product $x_{0}y_{0}$, and tracked their trajectories during GBE (Figs. S8B, C). The trajectories of the cells with the same initial coordinate product display hyperbolic characteristics, thus, cell center of gravity motions are those of an incompressible material undergoing convergent-extension flow.

The difference in the 2 families of cell trajectories is that, while both reflect incompressible flow, those in the anterior GB are fluid-like while those in the posterior GB are characteristic of a solid (Fig 7C). Posterior cell trajectories are more cohesive, with smoother and more collective movement patterns, suggesting that cells in this region translate together as a unified structure. In contrast, anterior cell trajectories exhibit greater fluctuations and a more dispersed pattern, indicating that cells in the anterior region are more likely to pass one another independently.

**Posterior pulling force penetration**

For simplicity, the posterior GB is assumed to behave as a purely elastic solid, and each row of cells parallel to the AP direction is considered identical. Therefore, an AP row of cells can be modeled as $N$ springs connected through $N+1$ beads. The $n$th spring connects bead $n-1$ and $n$. At $n=N$, the row of springs is subjected to a constant pulling force $F_{\mathrm{pull}}$, while at $n=0$ the boundary is stress-free (Fig. S9). The rest length of each spring is $b$. The position and velocity of the $n$th bead are denoted by $x_{n}$ and $v_{n}$, respectively, and the length of spring $n$ is $L_{n}$. Due to the pulling force applied to the $N$th bead, the springs are stretched and under tension, $T_{n}$, given by

$T_{n}=k\left( x_{n}-x_{n-1}-b \right)$,

(S8)

where $k$ is the one-dimensional effective spring constant. In our simulation, since the areal elasticity and the perimeter elasticity contribute similarly to the elastic energy, $k$ is taken to represent the areal elasticity. Accordingly, $\Delta E=1/2K_{a}\left( \Delta L\sqrt{A_{0}} \right)^{2}$, and hence $k=K_{a}A_{0}$.

As the $n$th bead connects both spring $n$ and spring $n+1$, force balance at the $n$th bead gives

$T_{n+1}-T_{n}-\lambda v_{n}=0$,

(S9)

where $\lambda$ is the drag coefficient representing friction between the cell apical surface and the overlying vitelline membrane. Substituting Eq. (S8) into Eq. (S9) yields

$k\left( L_{n+1}-b \right)-k\left( L_{n}-b \right)=\lambda v_{n}$,

$k\left( x_{n+1}-x_{n}-b \right)-k\left( x_{n}-x_{n-1}-b \right)=\lambda v_{n}$.

(S10)

Taking the continuum limit in the AP direction ($n\ll N$) and rewriting $v_{n}$ as $\dot{x}_{n}$ gives

$\dot{x}_{n}=\frac{1}{t_{0}}\frac{\partial^{2}x_{n}}{\partial n^{2}}$,

(S11)

where $t_{0}=\lambda/k=\lambda/{K_{a}A_{0}}$ defines the relaxation time per spring. Thus, relaxation time of the entire tissue is $t_{\mathrm{relax}}=N^{2}t_{0}$.

From dimensional analysis of Eq. (S11),

$\frac{\Delta x_{n}}{t}\sim\frac{1}{t_{0}}\frac{\Delta x_{n}}{n^{2}}$,

which gives

$n\sim\left( \frac{t}{t_{0}} \right)^{1/2}=\left( \frac{K_{a}A_{0}t}{\lambda} \right)^{1/2}$.

(S12)

At $t=30 \min$, the end of GBE rapid phase, the effects of the posterior pulling force penetrate ~ 30 cells, which is comparable to the length of the posterior GB (Fig. 6A).

Suppose at time $t$, the posterior pulling force has penetrated $\delta n$ cells, whose mean velocity is $v_{N}\left( t \right)$. The total drag experienced by these $\delta n$ cells is therefore $\lambda\delta n$. Hence,

$\lambda\delta n\cdot v_{N}\left( t \right)\approx F_{\mathrm{pull}}$,

and

$v_{N}\left( t \right)\approx\frac{F_{\mathrm{pull}}}{\lambda\left( \frac{t}{t_{0}} \right)^{1/2}}$,

(S13)

as $\delta n\sim\left( t/{t_{0}} \right)^{1/2}$. At $t=30 \min$, $v_{N}\approx3 \mu m/min$, consistent with our simulation results (Fig. 5E).

**Calculating Cell Alignment,** $\boldsymbol{Q}$

We computed the cell alignment, $Q$, using the same protocol as in Ref.(7), enabling direct comparison with previously reported solid-to-fluid transitions.

A triangular tiling was constructed from the centers of mass of the cellular polygons. Each vertex of the cellular network gives rise to $M$ triangles, as the vertex belongs to $M$ cells, where $M\leq3$. For each such triangle, one corner is defined as the average position of the centers of mass of all $M$ associated cells, while the remaining two corners correspond to the centers of mass of two adjacent cells.

For each triangle, we computed a symmetric, traceless tensor $\mathbf{q}$ that quantifies triangle elongation. This tensor is derived from a shape tensor $\mathbf{s}$, which describes the affine deformation mapping an equilateral reference triangle onto the observed triangle. Let the corners of a given triangle

$m$ be located at positions $\mathbf{r}^{A}$, $\mathbf{r}^{B}$, $\mathbf{r}^{C}$, ordered counterclockwise. The shape tensor $\mathbf{s}$ is defined as

$\mathbf{s}=\left( \begin{matrix} r_{x}^{B}-r_{x}^{A} & r_{x}^{C}-r_{x}^{A} \\ r_{y}^{B}-r_{y}^{A} & r_{y}^{C}-r_{y}^{A} \end{matrix} \right)\left( \begin{matrix} 1 & 1/2 \\ 0 & \sqrt{3}/2 \end{matrix} \right)^{-1}$.

(S14)

The shape tensor $\mathbf{s}$ is decomposed into three contributions: an isotropic (trace) part $\mathbf{t}$, a symmetric traceless part $\tilde{\mathbf{s}}$, and an antisymmetric part $\mathbf{s}^{\alpha}$,

$\mathbf{s}=\mathbf{t}+\tilde{\mathbf{s}}+\mathbf{s}^{\alpha}$.

(S15)

These components are given by

$\mathbf{t}=\frac{1}{2}\mathrm{Tr}\left( \mathbf{s} \right)$,

(S16)

$\tilde{\boldsymbol{s}}=\frac{1}{2}\left( \mathbf{s}+\mathbf{s}^{T} \right)-\frac{1}{2}\mathrm{Tr}\left( \mathbf{s} \right)\mathbf{I}$,

(S17)

$\mathbf{s}^{\alpha}=\frac{1}{2}\left( \mathbf{s}-\mathbf{s}^{T} \right)$.

(S18)

The triangle rotation angle $\theta$ is then extracted from the antisymmetric and trace components according to

$\binom{\cos\theta}{\sin\theta}=a\binom{t_{xx}}{s_{xy}^{\alpha}}$,

(S19)

where $a$ is a normalization prefactor.

The triangle elongation tensor $\mathbf{q}$ is computed as

$\mathbf{q}=\frac{1}{\left| \tilde{\mathbf{s}} \right|}\sinh^{-1} \left( \frac{\left| \tilde{\mathbf{s}} \right|}{\left( \det\left( \mathbf{s} \right) \right)^{1/2}} \right)\tilde{\mathbf{s}} \mathbf{R}\left( -\theta\right)$,

(S20)

where

$\left| \tilde{\mathbf{s}} \right|=\left( {\tilde{s}_{xx}}^{2}+{\tilde{s}_{xy}}^{2} \right)^{1/2}$,

(S21)

$\det\left( \mathbf{s} \right)$ is the determinant of the shape tensor $\mathbf{s}$, and $\mathbf{R}\left( -\theta\right)$ is a clockwise rotation matrix defined as

$\mathbf{R}\left( -\theta\right)=\left( \begin{matrix} \cos\theta& \sin\theta\\ -\sin\theta& \cos\theta\end{matrix} \right)$.

(S22)

The cell shape alignment tensor $\mathbf{Q}$ is obtained by averaging the triangle elongation tensors $\mathbf{q}_{m}$ over all triangles, weighted by their areas $a_{m}$,

$\mathbf{Q}=\left\langle\mathbf{q} \right\rangle$,

(S23)

with

$\left\langle\mathbf{q} \right\rangle=\frac{\sum_{m} a_{m}\mathbf{q}_{m}}{\sum_{m} a_{m}}$.

(S24)

Finally, the scalar cell shape alignment parameter is defined as the magnitude of $\mathbf{Q}$,

$Q=\left( {Q_{xx}}^{2}+{Q_{xy}}^{2} \right)^{1/2}$.

(S25)

**Figures**

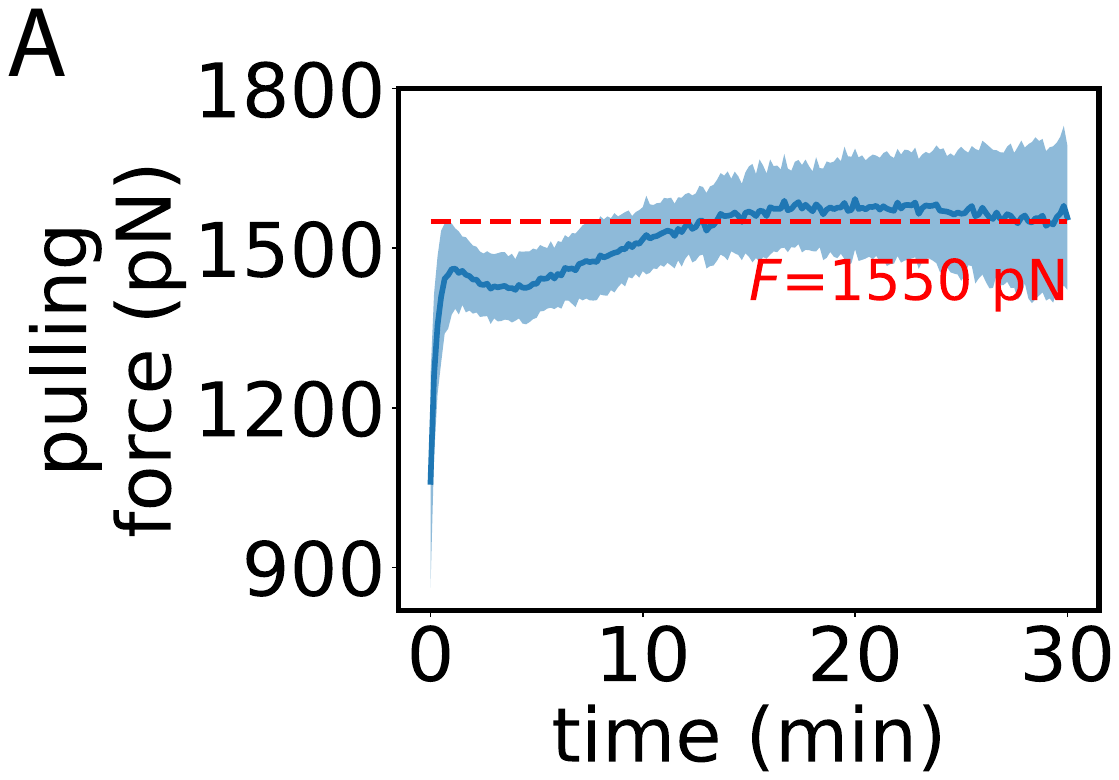

**Figure S1. Posterior pulling force as a function of time.**

(A) Posterior pulling force required to maintain constant velocity of the posterior vertices along the AP axis ($n=10 \mathrm{embryos}$). The force exerted on each vertex is calculated based on the force balance described in Eq. S1, Eq. S2. Vertex forces are then redistributed to cells according to vertex–cell associations. Shaded region denotes s.d..

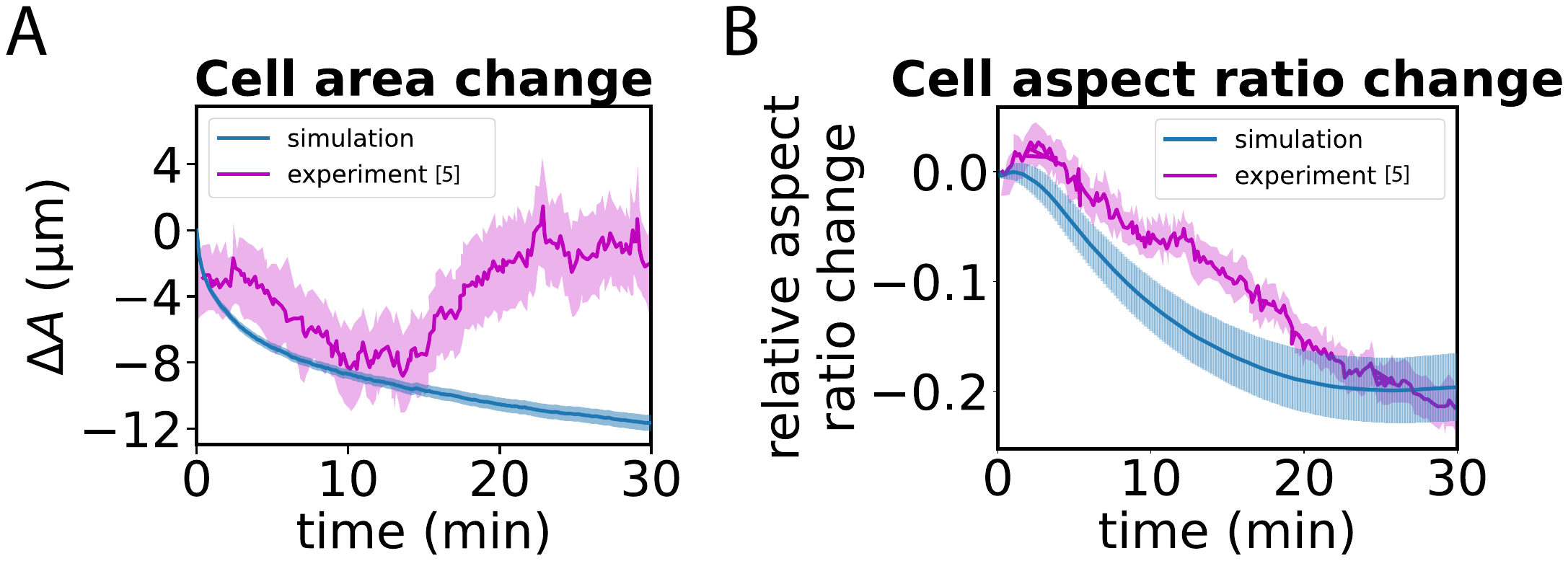

**Figure S2. Cell area change and cell aspect ratio change in simulation and experiment.**

(A) Cell area changes as a function of time in simulation and experiment. (B) Relative cell aspect ratio changes in simulation and experiment. Bars (simulation) and shaded region (experiment) represents s.d.. Experiment is replotted from ref (5).

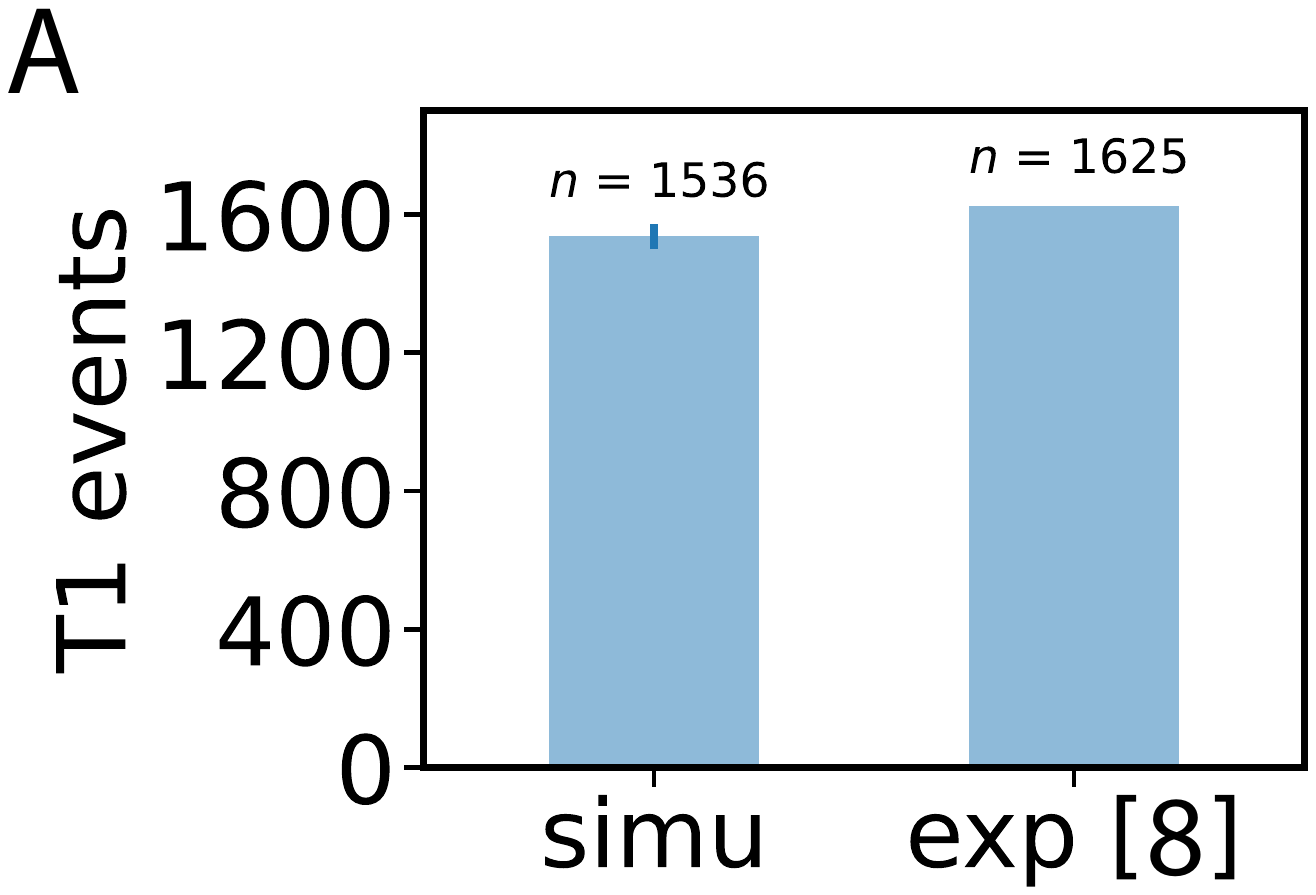

**Figure S3.** Total number of T1 transitions in WT time simulation (simu) and experiment (8) (exp). Bar represents the s.d..

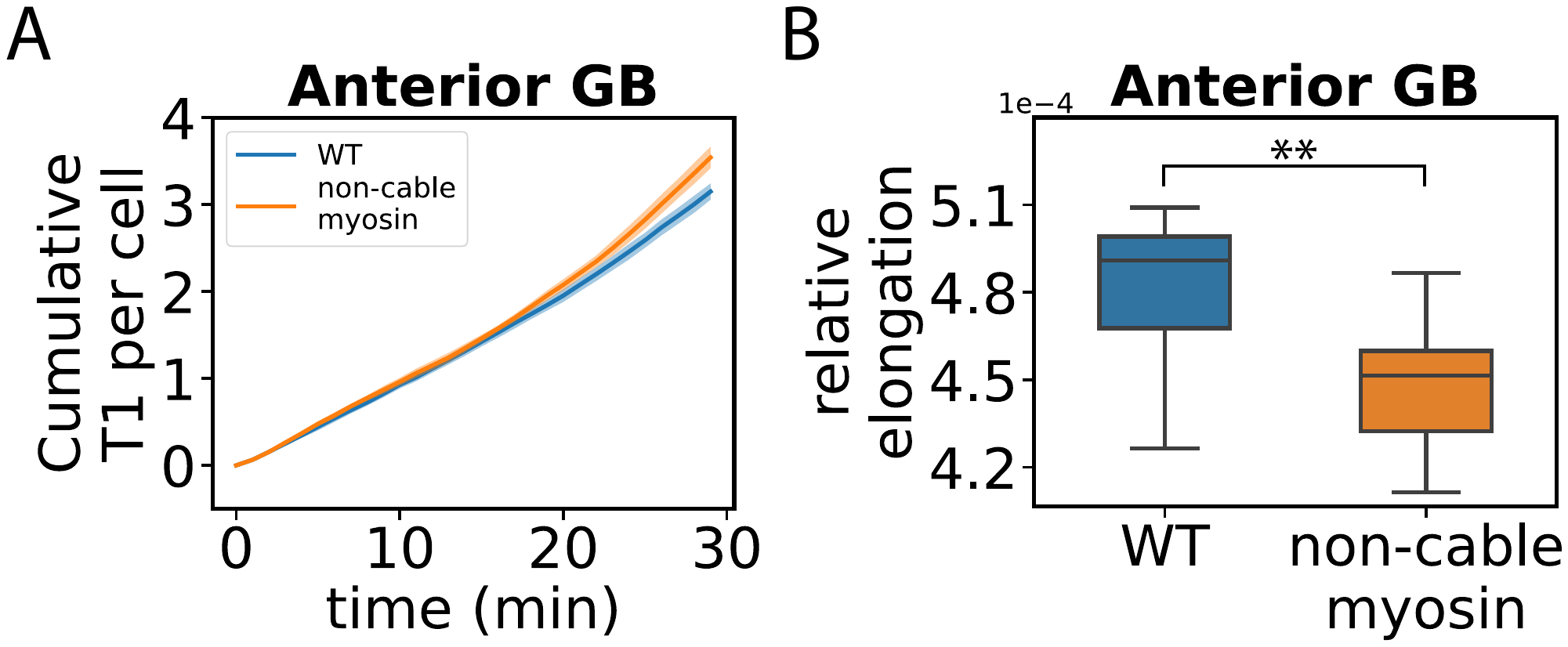

**Figure S4. T1 transitions have higher efficiencies when organized by myosin cables.**

(A) Cumulative T1 transitions per cell over time for the WT ($n=10 \mathrm{embryos}$) and the non-cable myosin mutant ($n=10 \mathrm{embryos}$). Shaded areas represent the s.d.. (B) Boxplot showing the distribution of the relative anterior GB elongation contributed by individual T1 events in the WT and the non-cable myosin mutant ($n=10 \mathrm{embryos}$ for both). Significance indicators used here and throughout the paper: NS, p > 0.05; * p < 0.05; ** p < 0.01; *** p < 0.001.

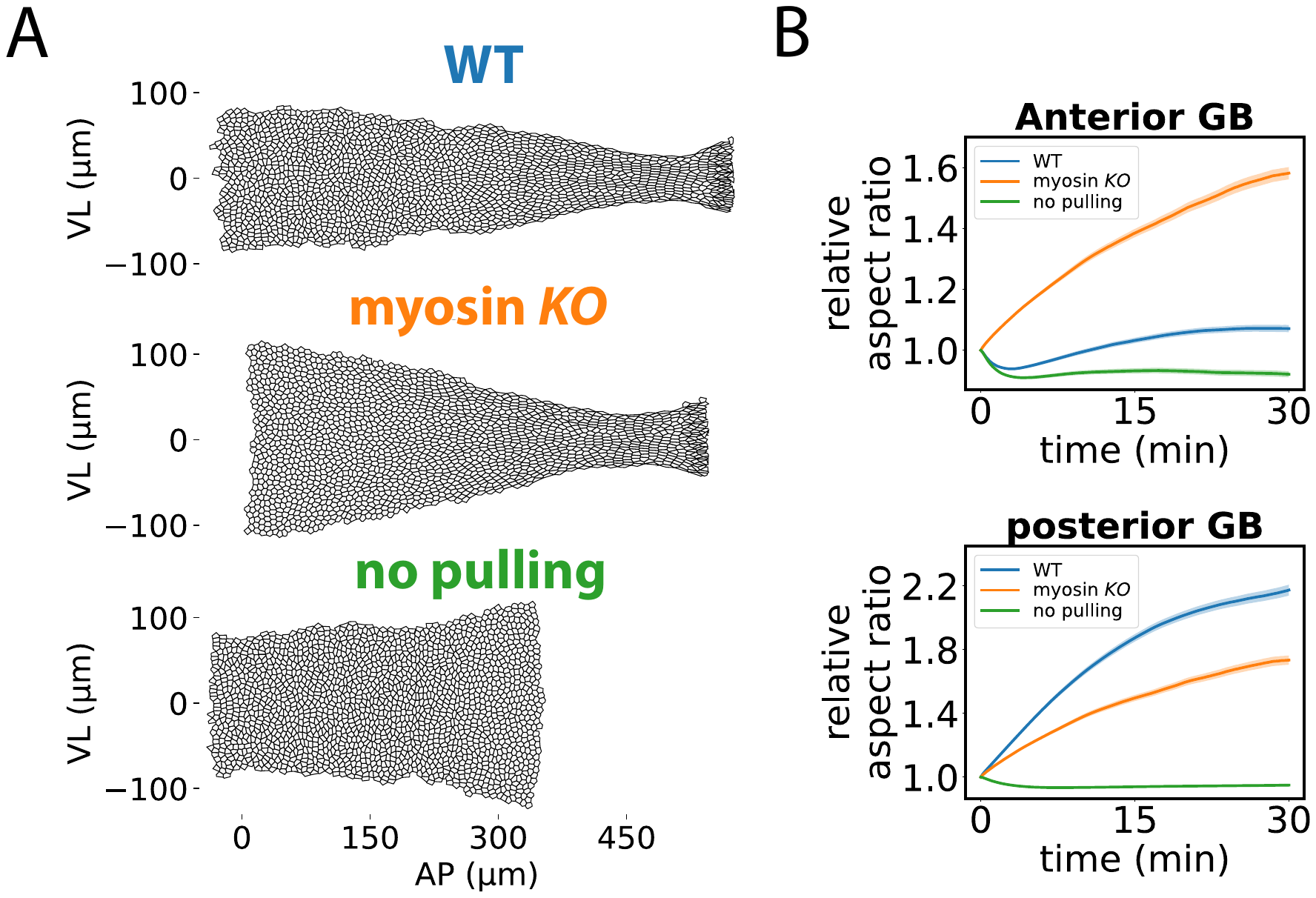

**Figure S5. Distinct germ-band shapes and cell morphologies in the WT and mutant embryos**

(A) Example outcome shapes of the germ band at $t=30 \min$ in the WT, myosin *KO* mutant, and mutant lacking posterior pulling force. Corresponding cell morphologies are shown. (B) Cell aspect ratio as a function of time for the anterior GB (top) and the posterior GB (bottom) in the WT, myosin *KO* mutant, and no-pulling-force mutant ($n=10 \mathrm{embryos}, 1853 \mathrm{cells}$ for each). Shaded areas represent the SEM.

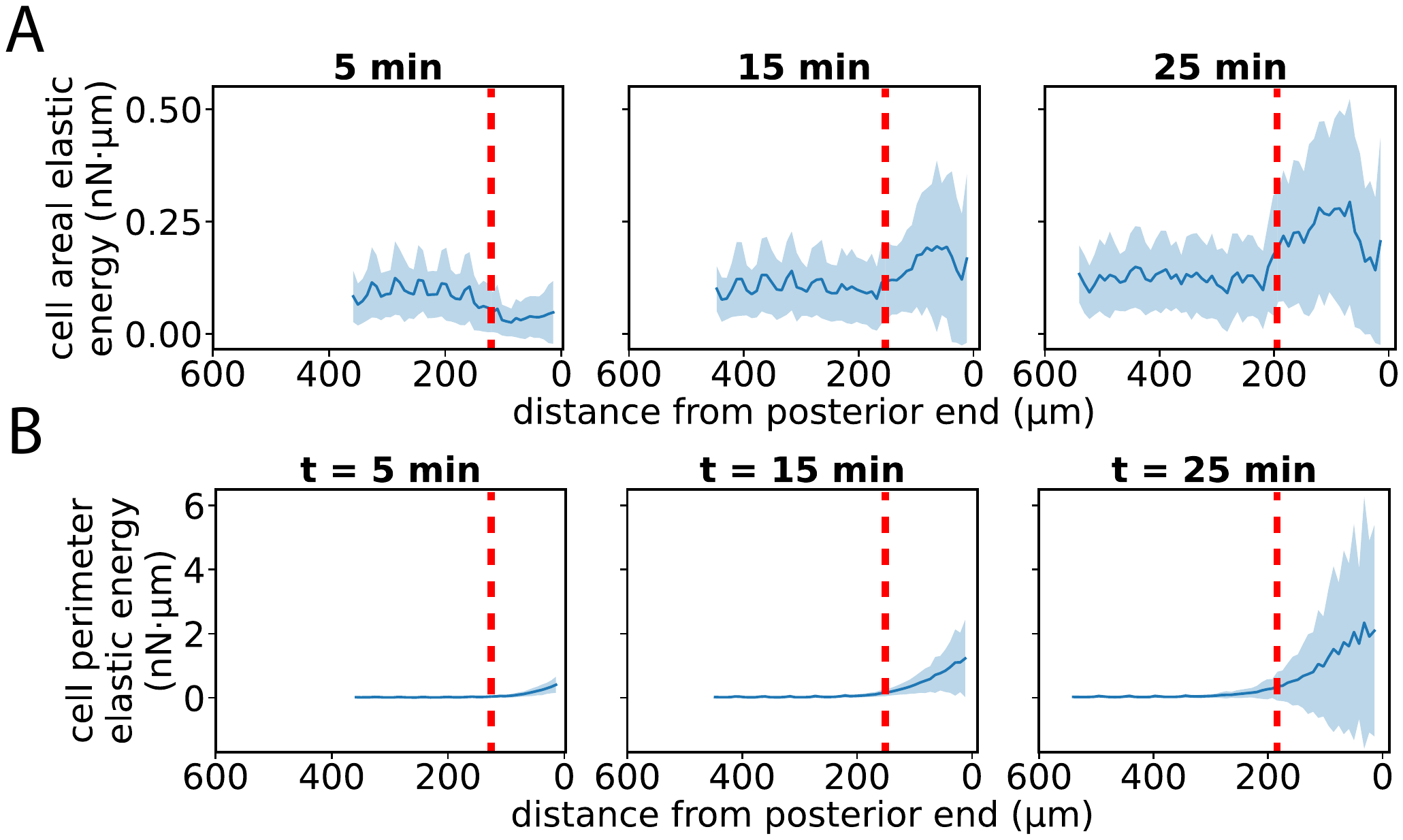

**Figure S6. Cell areal and perimeter elastic energy**

Cell areal (A) and perimeter (B) elastic energy versus location in the AP direction at the indicated times during GBE ($n=10 embryos, 1853 \mathrm{cells}$ each). Red dashed line indicates the boundary between the anterior GB and the posterior GB in (A) and (B). Shaded regions for both (A) and (B) represent s.d..

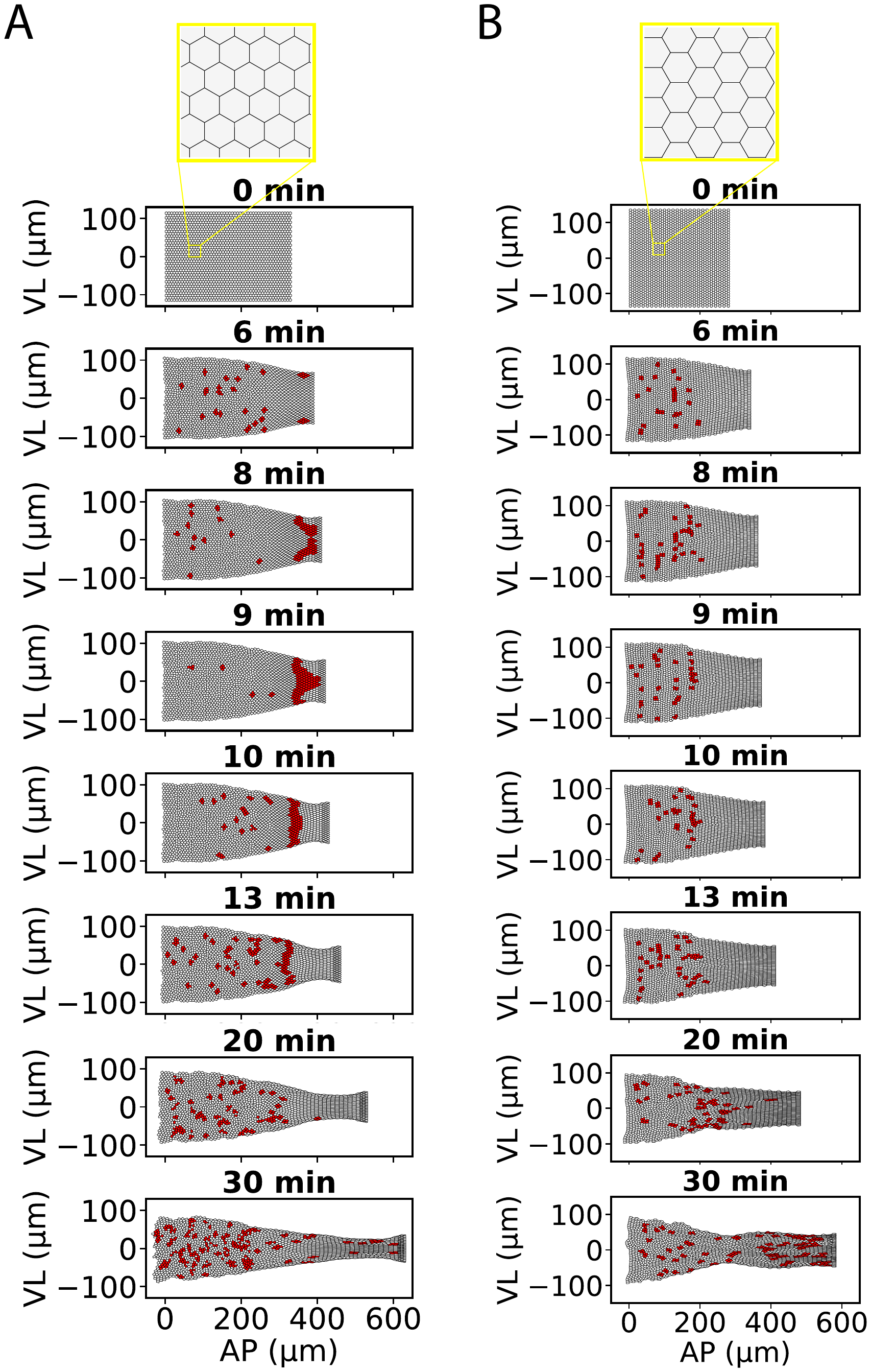

**Figure S7. Distinct patterns of T1 transitions in perfect-lattice initial conditions with different orientations.** (A) VL-oriented perfect hexagonal lattice, in which cells of the initial lattice have two edges parallel to the VL axis. (B) AP-oriented perfect hexagonal lattice, in which cells of the initial lattice have two edges parallel to the AP axis. Cells undergoing T1 transitions within 1 min of each selected time point are marked in red.
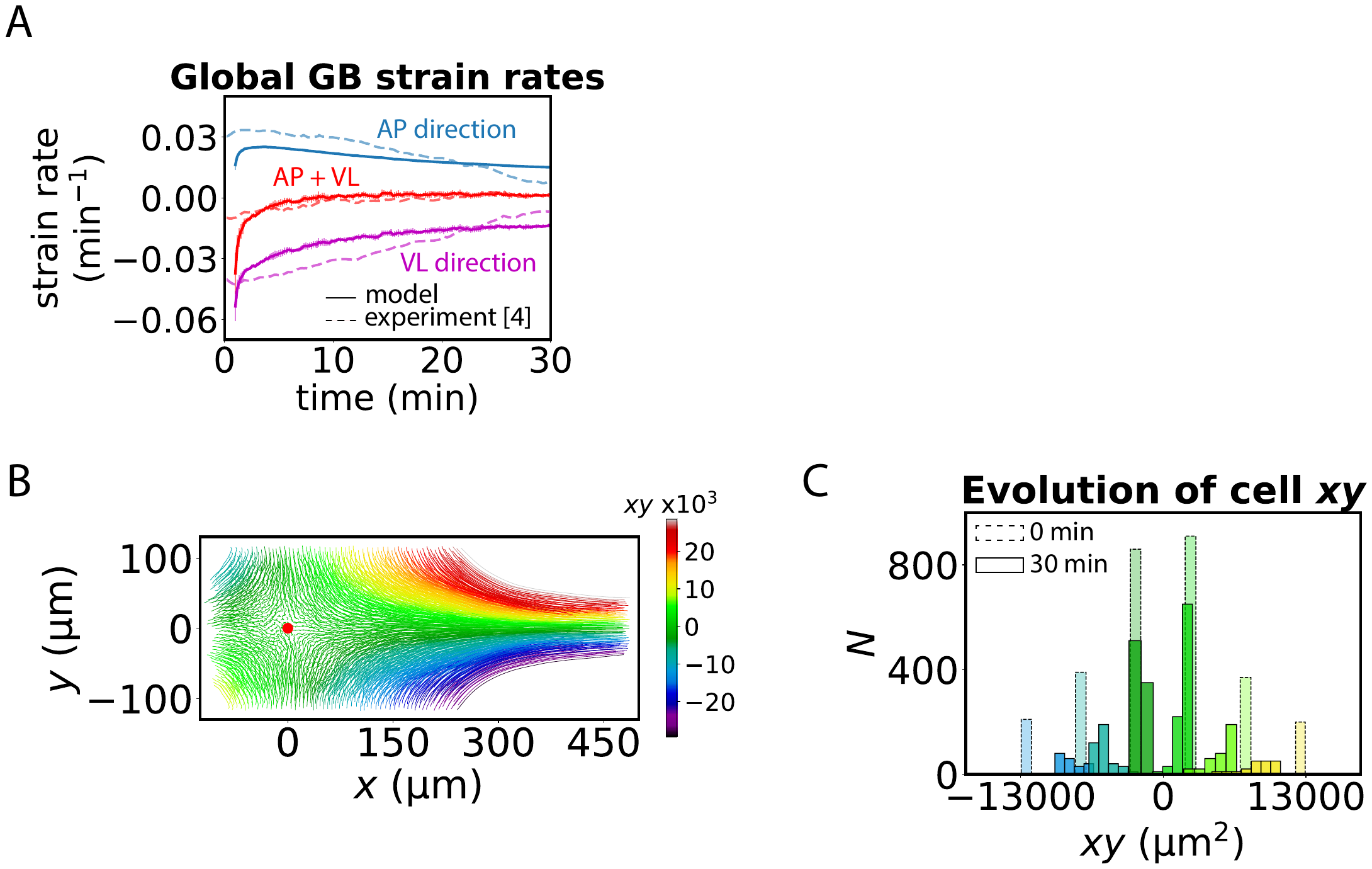

**Figure S8. Both anterior and posterior GB cell trajectories are hyperbolic.** (A) Global GB strain rates versus time in the AP and VL directions predicted by model ($N=10$ embryos) and observed experimentally ($N=5$ embryos). Model AP strain rate defined as rate of change of mean relative locations of anterior and posterior germ-band boundaries in AP direction, averaged over boundary vertices; similarly for VL strain. Throughout GBE the tissue flow is approximately incompressible, since the sum of the strains (red curves) is small compared to the strains. Experimental data replotted from ref (4). Errors (curve thicknesses) are s.d.. (B) Model-predicted trajectories of all cells during the 30 min of GBE. The origin of coordinates (red disc) is chosen as the location of zero velocity. (C) Histogram of coordinate products $x\left( T \right)y\left( T \right)$ at the end of GBE, $T=30$ min (solid columns) for several values of the initial product $x_{0}y_{0}$ (dashed columns). Color scheme identifies cells with the same initial coordinate product $x_{0}y_{0}$ (within $\pm500\text{ }\text{μm}^{\text{2}}$) in both (B) and (C). $x$ and $y$ represent the AP and VL positions of the cells relative to the trajectory origin ($\mathrm{AP}=127 \mu m, VL=0 \mu m$).

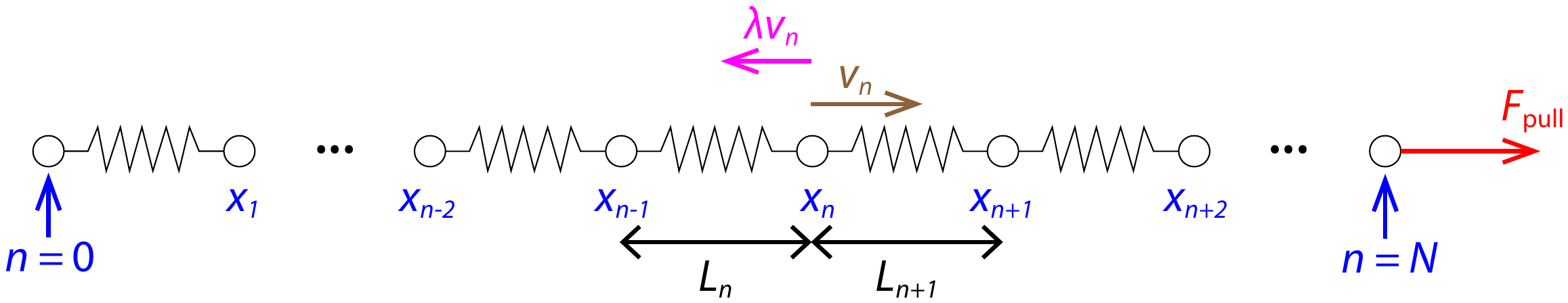

**Figure S9. Schematic of the simplified germ band (GB) represented in one dimension.** Cells are modeled as springs, with a total of N cells. White discs, equivalent to cell–cell junctions in 2D, represent beads that connect neighboring springs (cells). Each spring $n$ is associated with two beads, $n-1$ and $n$, whose positions and velocities are denoted by $x_{n}$ and $v_{n}$, respectively. $L_{n}$ indicates the length of the $n$th spring. Brown arrow, bead velocity in response to the pulling force. Red arrow, pulling force exerted on the boundary at $n=N$. Magenta arrows, drag forces from the vitelline membrane acting opposite to bead velocities. Black double arrows, lengths of the corresponding springs.

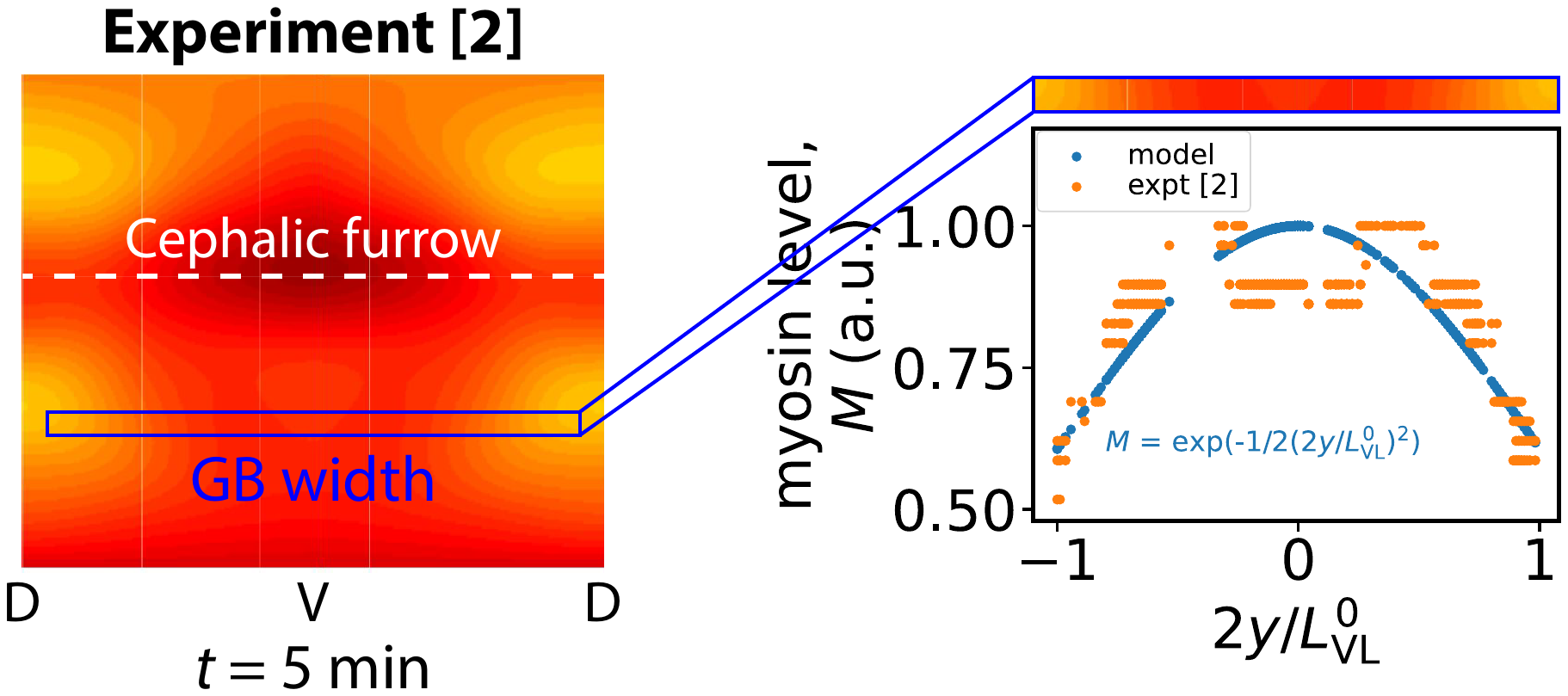

Figure S10. Myosin intensity exhibits a bell-shaped profile in the VL direction. Apical myosin level measured in experiment from ref. (2) (left). The blue box marks the region used to extract the spatial envelope; its width approximately corresponds to the germ-band (GB) width. Myosin intensities in the experimental heat map were rescaled from 0 to 1, where 0 and 1 represent the minimum and maximum signal levels, respectively. In the experimental image, dark red indicates high myosin intensity, and bright yellow indicates low intensity. The curve labeled $M$ (right) shows the best-fit function describing the extracted myosin profile.
